# Land-use impacts on plant functional diversity throughout Europe

**DOI:** 10.1101/2024.02.21.580945

**Authors:** Francesca Rosa, Peter M. van Bodegom, Stefanie Hellweg, Stephan Pfister, Idoia Biurrun, Steffen Boch, Milan Chytrý, Renata Ćušterevska, Michele Dalle Fratte, Gabriella Damasceno, Emmanuel Garbolino, Jonathan Lenoir, Wim A Ozinga, Josep Penuelas, Francesco Maria Sabatini, Franziska Schrodt, Domas Uogintas, Chaeho Byun, Jiri Dolezal, Tetiana Dziuba, Bruno Hérault, Irene Martín-Forés, Ülo Niinemets, Gwendolyn Peyre, Laura Scherer

## Abstract

**Aim:** Global biodiversity loss resulting from anthropogenic land-use activities is a pressing concern, requiring precise assessments of impacts at large spatial extents. Existing models mainly focus on species richness and abundance, lacking insights into ecological mechanisms and species’ roles in ecosystem functioning. To bridge this gap, we conducted an extensive analysis of the impact of human land use on vascular plant functional diversity, across diverse land-use classes and bioregions in Europe, comparing it to traditional metrics.

**Location**: Europe **Time period**: 1992-2019 **Major taxa studied**: Vascular plants

**Methods:** Integrating extensive databases of vegetation plots with spatial data on land use and land cover, we paired plots from areas actively used and modified by humans with plots from natural habitats under similar environmental conditions. Using species occurrences and traits, in each plot we computed three complementary functional diversity metrics (functional richness, evenness, and divergence), species richness and abundance. We assessed the impact of land use by comparing the metrics in the paired plots.

**Results:** Our findings revealed that, compared to natural habitats, anthropogenic land use exhibits lower functional richness and divergence but higher functional evenness across most land-use classes and bioregions. The response of functional richness was more marked than the other two metrics and especially pronounced in croplands and urban areas and in northern bioregions. Functional richness exhibited a pattern that did not fully overlap with the trend in species richness, providing useful complementary information.

**Main conclusions:** We provide a large-scale precise assessment of anthropogenic land-use impacts on functional diversity across Europe. Our findings indicate that: (i) human disturbance significantly alters plant functional diversity compared to natural habitats; (ii) this alteration goes in the direction of functional homogenization; (i) functional diversity metrics complement traditional metrics by offering deeper insights into the ecological mechanisms in response to anthropogenic land use.

## 1. Introduction

Biodiversity plays a crucial role in bolstering ecosystem functions, maintaining services vital for all organisms, including humans, and mitigating some global changes (Isbell et al. 2011; Harrison et al. 2014; Tilman et al. 2014; Felipe-Lucia et al. 2018; Valladares et al. 2019; Le Provost et al. 2023; Reader et al. 2023). Anthropogenic activities, particularly habitat loss and degradation, pose significant threats to biodiversity (IPBES 2019; Pereira et al. 2024). With over 80% of global land transformed due to human actions (Ellis et al. 2021)e, ecosystems and their functions face severe impacts, underlining the urgent need for comprehensive strategies to halt biodiversity loss (Leclère et al. 2020; Carmona et al. 2021) and for a better understanding of the linkage between drivers and effects at different spatial scales (Chaplin-Kramer et al. 2022).

To effectively tackle this challenge, it is essential to expand our comprehension of biodiversity beyond conventional metrics like species richness or abundance, which are frequently employed in current biodiversity models given their availability (O’Connor et al. 2017; Bannar-Martin et al. 2018; Jandt et al. 2022). However, these metrics often fail to capture the ecological significance of species within a community or their role in ecosystem functioning (Suárez-Castro et al. 2022) and exhibit considerable variability in trends (Dornelas et al. 2023; Valdez et al. 2023). Therefore, it is imperative to explore additional dimensions of biodiversity, such as functional diversity, here defined as the variation and distribution of species’ functional traits within communities. Functional diversity enhances and complements traditional metrics by providing deeper knowledge into ecological mechanisms through the integration of functional information (Cadotte et al. 2011; Hu et al. 2014; Rosenfield and Müller 2020; Scherer et al. 2023). Particularly, the functional diversity of plants has been shown to offer further insights into ecosystem performance than taxonomic diversity (Bruelheide et al. 2018; van der Plas 2019; Zambrano et al. 2019; Kattge et al. 2020; Hagan et al. 2023), especially in areas affected by human activities (Bonilla-Valencia et al. 2022).

To calculate functional diversity, three independent and complementary indices are commonly used: functional richness, evenness, and divergence (Mason et al. 2005). These indices, derived from traits encompassing anatomical, physiological, biochemical, regenerative, reproductive, and phenological characteristics, provide valuable insights into ecosystem dynamics (Villéger et al. 2008). Functional richness represents the amount of functional niche space filled by a species assemblage. Functional evenness describes how regularly species abundances are distributed in the functional niche space. Functional divergence measures to which degree species abundances are distributed from the centre of the functional space to its extremes; it is sensitive to the presence and abundance of species with extreme trait values, e.g., highly specialized or functionally rare species (Mason et al. 2005; Villéger et al. 2008; Mouchet et al. 2010; Scherer et al. 2020).

Along with the need of complementary biodiversity metrics, currently there is a lack of well-established connections between individual local biodiversity assessments and global patterns (Jandt et al. 2022), hindering a comprehensive analysis and suggesting the need for replicated local assessments (Knollová et al. 2024). These assessments are particularly crucial in understanding the effect of human use of land, which encompasses activities like agriculture and urbanization (hereafter simply referred to as “land use”), compared to natural habitats (Zebisch et al. 2004; Dornelas et al. 2014; Jandt et al. 2022).

Previous studies have explored the effect of land use on plant functional diversity at a large spatial extent, from national to global (Flynn et al. 2009; De Souza et al. 2013; Carmona et al. 2020; Newbold et al. 2020; Scherer et al. 2020). However, these studies often used different and less comprehensive indices, failed to distinguish impacts across regions, only focused on a specific land-use class or did not compare functional diversity with more common metrics.

In this research work, we leverage the recent release of global vegetation and trait databases to explore the effect of land use on different dimensions of functional diversity, especially in regions with good data coverage and representativeness, such as Europe (Chytrý et al. 2016; Bruelheide et al. 2019; Kattge et al. 2020). We utilized data from the European Vegetation Archive (EVA) (Chytrý et al. 2016), a vegetation plot database containing information on species co-occurrences and abundances within plant communities. Coupled with the TRY database, which provides species-level plant trait data (Kattge et al. 2020), it allows for the calculation of functional diversity for approximately two millions of vegetation plots across all of Europe (Bruelheide et al. 2018). By matching vegetation plots and trait data with land use and land cover, we investigated the change in functional diversity in anthropogenic land compared to natural and close-to-natural habitats across Europe.

Given this context, our goal was to answer the following research questions: How does plant functional diversity change in land actively used by humans compared to natural habitats? Which additional information does functional diversity provide compared to other metrics? Are the biogeographic conditions affecting the response?

## 2. Materials and methods

We compared the functional diversity in vegetation plots located in anthropogenic and natural land by combining species and trait information with land-use and land-cover maps.

We applied seven steps (see Fig. 1 and, for more details, Fig. S1.1). First, we retrieved and filtered the suitable vegetation plots from the European Vegetation Archive (EVA) (Chytrý et al. 2016), while matching them to the sPlot database (Bruelheide et al. 2019), which contains all of the EVA plots and has curated a taxonomic standardization procedure to link each species name to a set of 30 gap-filled traits from the TRY database (Schrodt et al. 2015; Kattge et al. 2020) (see section 2.1). Second, we assigned each vegetation plot to a biogeographic region (hereafter “bioregion”) to enable a spatially explicit analysis while including a sufficient sample size per spatial unit (see section 2.2). Third, based on land-use and potential natural vegetation maps, we aggregated land-use classes into broader ones suitable for the analysis (see section 2.3). Fourth, we assigned each vegetation plot to one of the five identified anthropogenic land-use classes (*Urban areas*, *Cropland*, *Pasture and rangeland*, *Mosaic*, *Planted forest*) or to one of the five identified potential natural vegetation classes (*Natural forest*, *Natural grassland*, *Natural shrubland*, *Natural herbaceous wetland*, *Natural bare and sparse vegetation*) (see section 2.4). Fifth, we selected the relevant environmental variables (i.e., bioclimatic variables and soil properties) and then performed a Principal Component Analysis (PCA) of these variables across Europe to obtain the scores of each vegetation plot along the PCA axes (see section 2.5). Sixth, to allow for a consistent comparison between anthropogenic land-use classes and natural habitats, we paired each vegetation plot sampled in anthropogenic land with a vegetation plot sampled in the natural habitat that would potentially occur there under low human pressure. To do that, we selected the pairs from the same bioregion with minimal distance of PC scores in the PCA space (see section 2.6). Finally, we computed functional richness, functional evenness, functional divergence, species richness, and total abundance in each plot; we then calculated the relative values of these metrics in each vegetation plot sampled in anthropogenic land compared to the paired vegetation plot in natural habitat (see section 2.7).

**Figure 1.**
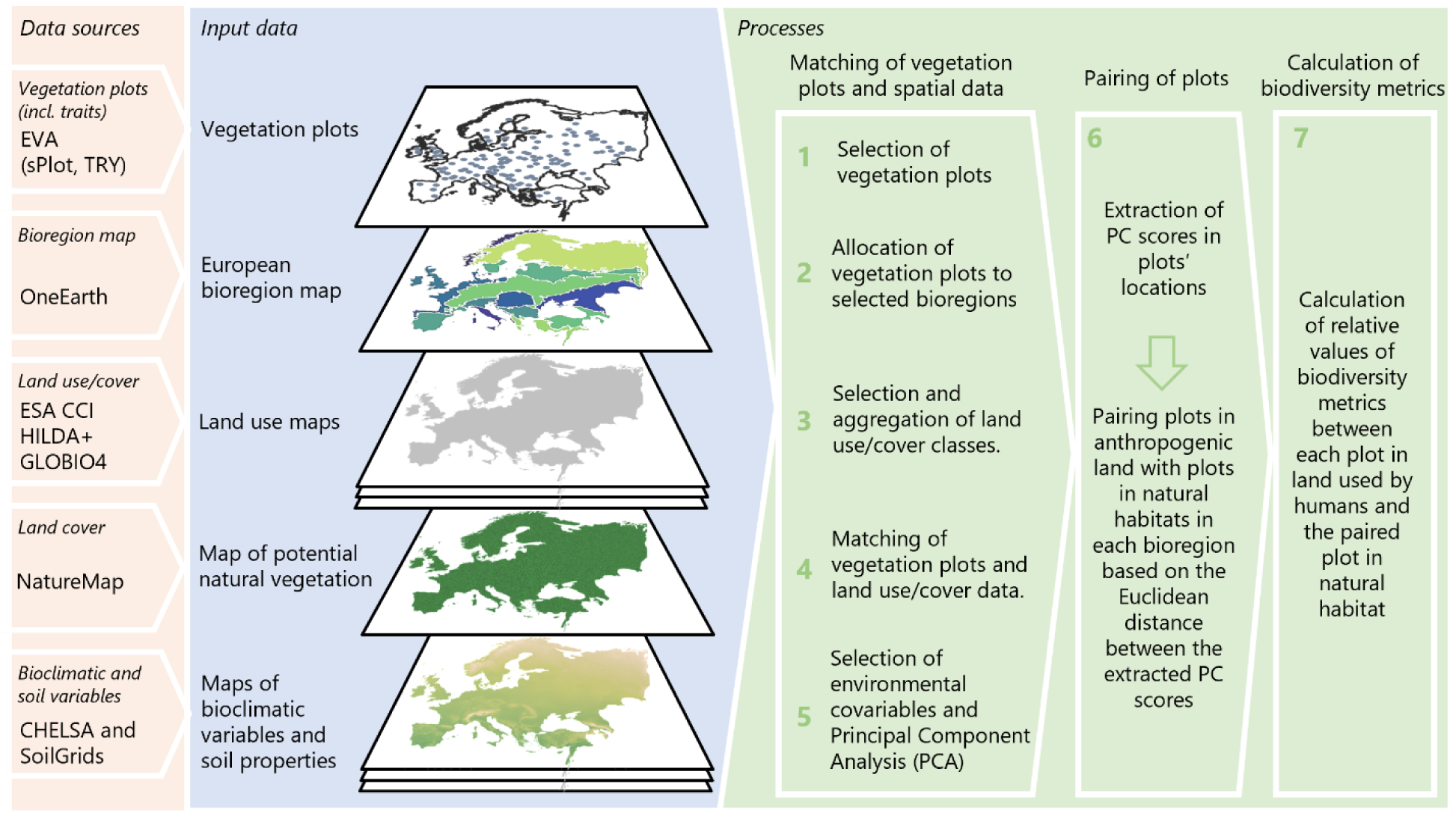
Overview of the methodological conceptual framework applied to this study.

All analyses were performed in R version 4.0.5 (2021-03-31) (R Core Team 2021).

### 2.1 Selection and processing of vegetation plots

We extracted vegetation plot data from EVA (Chytrý et al. 2016). It currently contains more than 2 million vegetation plots and has a representative geographical coverage for Europe, especially for West, South and Central Europe. We used data on 30 gap-filled plant traits from TRY (Schrodt et al. 2015; Kattge et al. 2020) and an additional plot attribute (plot naturalness, used in section 2.4) from sPlot (Bruelheide et al. 2019). Not all the vegetation plots were suitable for the analysis, as some do not report, for example, geographical coordinates and/or the year of the survey. Therefore, we applied various filtering criteria (see section S2 for the full list) and selected a subset of datasets from EVA (see Table S16 for the list).

We considered only those vegetation plots for which species abundance data were available, as this information was needed to calculate functional evenness and divergence. Furthermore, we retained only vascular plant species with available traits. Additional vegetation plots were excluded from the analysis for various reasons: e.g., when trait information was known for a small proportion (< 0.5) of species occurring in the plot, or the location uncertainty was too high (Engel et al. 2023).

We set a threshold for location uncertainty of 10 km, because a stricter threshold would exclude almost all plots from some regions (e.g., the Iberian Peninsula). Since this threshold is considerably higher than the resolution of the land-use maps (300 m, see section 2.3), for plots with location uncertainty higher than 150 m, we applied an additional filter based on the homogeneity of the land use of the area falling within the uncertainty radius of the plot. After matching land-use and land-cover classes to the vegetation plots (as explained in section 2.4), we retained only plots where at least 80% of the land use or land cover within the uncertainty radius was the same as that occurring at the location of the plot coordinates. A ranking was made to keep track of this procedure (ranking 1: location uncertainty < 150 m; ranking 2: location uncertainty > 150 m and land use 100% homogeneous within the uncertainty radius; ranking 3: location uncertainty > 150 m and land use being 80 to < 100% homogeneous within the uncertainty radius). The ranking was later used to refine the pairing between vegetation plots in anthropogenic land and vegetation plots in natural habitats (as explained in section 2.6).

### 2.2 Allocation to bioregions

To assess spatial variations in the effects on functional diversity, we assigned vegetation plots to bioregions (One Earth). This approach enabled regionalization, while facilitating an appropriate sampling size, as higher resolution of other more commonly used regionalizations (e.g., (Olson et al. 2001), would have drastically reduced the sample size (list of bioregions’ names available in Table S3.1 and map in Fig. 2 and S3.1).

**Figure 2.**
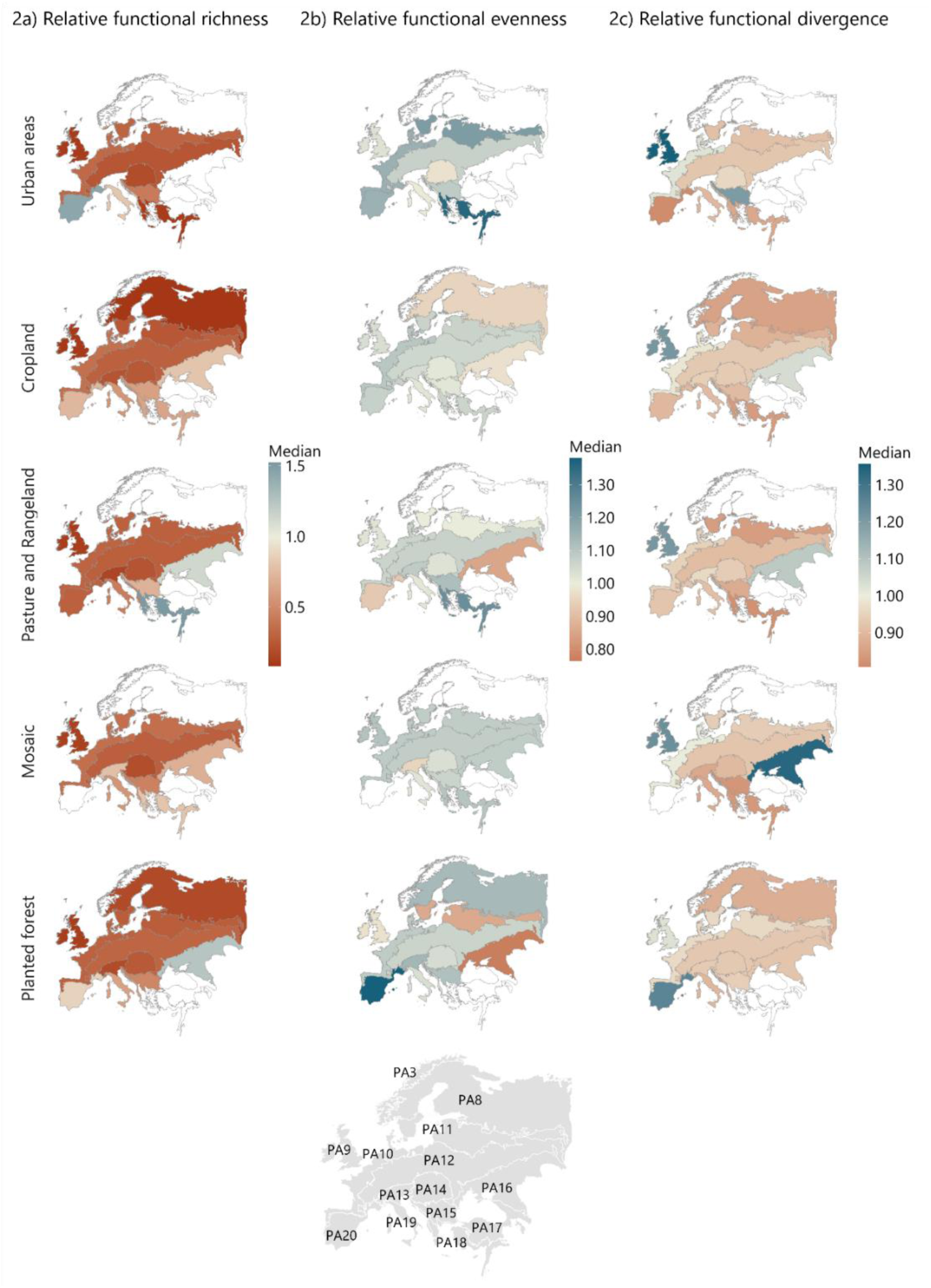
Median values of relative functional richness (a), evenness (b) and divergence (c) per bioregion and land-use class. Warmer colors and colder colors indicate decreased and increased functional diversity values, respectively, in relation to the potential natural vegetation. White bioregions are those for which not enough pairings were available.

### 2.3 Selection and processing of land use, land cover and potential natural vegetation maps

Each vegetation plot was matched to either an anthropogenic land-use class or to a class of natural vegetation (described in more detail in section 2.4).

As a base layer for land-use classes, we adopted the land-use and land-cover map from the European Space Agency for the Climate Change Initiative (ESA CCI) (ESA Land Cover CCI project team and Defourny 2019). The ESA CCI map had several advantages: (1) it has a 300 m spatial resolution, which we deemed adequate for our goals; (2) it includes most of the land uses and land covers we are addressing in this study (urban areas, cropland, natural vegetation), although not all (e.g., it does not distinguish between natural and managed grassland or forest); (3) it has a yearly resolution, although it goes back only to 1992 (so plots sampled before that year had to be excluded); and (4) it is open access. To fill the gaps concerning some land-use classes, we integrated the ESA CCI data with: (i) the HILDA+ map (Winkler et al. 2021), which covers the period 1960-2019 on a yearly basis and distinguishes between natural grass/shrubland and human-modified pasture/rangeland) and (ii) the GLOBIO4 map (Schipper et al. 2019; PBL Netherlands Environmental Assessment Agency 2023), which was built using the ESA CCI as a basis and distinguishes planted forests from the other land uses and land covers. GLOBIO4 provides data for 1992, 1995, 2000, 2005, 2010, 2015. The land use of the intermediate years was assumed to be the same as the closest year for which the map was available (e.g., for 1993, 1992’s data were used). Bioregions PA1 (Russian Arctic Desert Islands) and PA2 (Iceland) were excluded, as they did not contain any vegetation plot in anthropogenic land uses.

Concerning the map of potential natural vegetation, the NatureMap was selected (Hengl et al. 2020), as its resolution (250 m) is similar to the one of the ESA CCI map, and it allows for a distinction between multiple types of natural vegetation (e.g., forest, grassland, shrubland, etc.).

ESA CCI contains 38 land-use/cover classes and the NatureMap 17 land-cover classes. An aggregation was performed to reduce the number of classes and align the classification with the scope of the study (see Table S4.1 and Table S5.1 for the detailed matching between the aggregated classes and, respectively, the ESA CCI classes and the NatureMap classes).

### 2.4 Assigning vegetation plots to anthropogenic land-use classes or classes of natural habitat, and to the potential natural vegetation

Using the geographical coordinates of the center of each vegetation plot and its sampling date, we extracted the corresponding land-use and land-cover classes. The list of land-use and land-cover classes retrieved from the spatial sources described in the previous section and how they were combined to define the finally used classification (Table 1) is available in Table S6.1. Two additional attributes were used to refine the matching between plots and land-use/cover classes: (i) the type of vegetation from the EVA database (forest, shrubland, grassland, sparse vegetation or wetland); (ii) and the level of naturalness from the sPlot database (1: natural, 2: semi-natural, 3: anthropogenic) (sPlot 2024). In the final classification, each plot was assigned to either a anthropogenic land-use class or a natural habitat (Table 1).

**Table 1.** List and definition of the land-use/cover classes considered in this study, retrieved from the combination of spatial data, and assigned to the vegetation plots. The list of original land-use and land-cover classes and how they were combined is shown in Table S6.1.

| Land use/cover assigned to the plots |  | Definition |
| --- | --- | --- |
| Natural and close-to-natural habitats | Natural forest | Areas where vegetation belongs to the same formation and type that would occupy the site without human interference. This includes forests with sporadic wood extraction or natural grasslands that are grazed at such low intensity that the natural vegetation type is not replaced. |
|  | Natural grassland |  |
|  | Natural shrubland |  |
|  | Natural herbaceous wetland |  |
|  | Natural bare and sparse vegetation |  |
| Anthropogenic land-use classes | Planted forests | Planted, intensively managed forests. It excludes forest planted for protection, ecosystem restoration, or established through planting or natural regeneration, mimicking natural forest stands at maturity. |
|  | Pasture and rangeland | Herbaceous land or shrubland used for livestock and grazing with different intensities and practices. It excludes areas grazed at very low intensity (see two rows above). Vegetation plots assigned to *Pasture and rangeland* do not necessarily have *Natural grassland* as potential natural vegetation, because in most of Europe potential natural vegetation is *Natural forest*, even where now there are pastures and rangelands. |
|  | Cropland | Agricultural land where annual or permanent crops (trees, shrubs and herbaceous) are grown. |
|  | Urban areas | Human settlements, including artificial surfaces, built-up areas, industrial areas, and urban green spaces. |
|  | Mosaic | Areas characterized by a combination of natural vegetation and cropland to varying degrees. |

### 2.5 PCA on the environmental variables

We retrieved 19 bioclimatic variables – temperature- and precipitation-related – at 1-km resolution from the CHELSA V2.1 database (Karger et al. 2017; Karger et al. 2018), and a selection of soil properties that are less influenced by land management (clay mass fraction, silt mass fraction, sand mass fraction, and pH) at 250-m resolution from the SoilGrids database (ISRIC; Hengl et al. 2017) (see the full list in Table S7.1). The soil variables were spatially aggregated to match the resolution of the bioclimatic variables by calculating their mean in each 1-km square. The PCA was performed for these variables at 1-km resolution and on the whole area under assessment, similarly to what was done by (Sabatini et al. 2021). Then we extracted the scores along all the PCA axes for each vegetation plot’s location. This approach enabled us to assign each vegetation plot to a position in the PCA space defined by the environmental conditions (Wallis et al. 2021; Joswig et al. 2022), which was later used to pair the plots (see section 2.6).

The PCA was performed using the function rasterPCA from the package RStoolbox, which also allows for a standardization of the input data (see Fig. S7.1 for the results of the first four PCA axes).

### 2.6 Pairing of vegetation plots in anthropogenic land use and natural habitats

After extracting the PC scores, we paired each vegetation plot sampled in an area assigned to a land-use class with a vegetation plot sampled in the corresponding potential natural land and sharing similar environmental conditions. This approach paired vegetation plots that are close to each other in the environmental space of the PCA as well as belonging to the same bioregion, minimizing the Euclidean distance between the vegetation plots’ PC scores within a given pair (see Fig. S8.1) (Elmore and Richman 2001). To calculate the Euclidean distance, each PC score was weighted according to the explained variance of the corresponding PCA axis. Multiple pairings were possible for the same vegetation plot in a natural habitat if the plot had a minimum distance with multiple vegetation plots in human-used land (Fig. S8.1).

The opposite situation also occurred: the same vegetation plot from an anthropogenic land-use class was paired with multiple vegetation plots belonging to natural habitats if the environmental distance within the PCA space was the same. When this occurred, only the vegetation plot from natural habitats with the lowest location uncertainty ranking was considered. If multiple natural plots remained paired to the same plot even after filtering out those with higher location uncertainty ranking, they were kept, and the average of their biodiversity metrics (see next section) was calculated.

Any combinations of bioregion and land-use class with fewer than 10 pairings were removed from the subsequent analyses for statistical reasons, and this caused the exclusion of two additional bioregions not fulfilling this requirement: PA3 (Scandinavian Birch & Coastal Conifer Forests) and PA17 (Black Sea, Caucasus-Anatolian Mixed Forests & Steppe).

### 2.7 Calculation of relative functional and species diversity

We calculated functional richness, evenness and divergence for each vegetation plot using the dbFD function from the FD package in R (Villéger et al. 2008; Laliberté et al. 2010; Ahmed et al. 2019). All plant traits were standardized to zero mean and unit variance. As our aim was to assess the response of the overall ecosystem functioning and not of specific ecosystem functions, we considered all 30 traits available in sPlot (full list in Table S10.1) from the gap-filled TRY database, as using imputed data was shown to be robust compared to a reduced species set (Scherer et al. 2023).The traits ranged from morphological and nutritional to reproductive characteristics, including stem-specific density, rooting depth, specific leaf area, leaf carbon and nitrogen content, plant height, seed characteristics, leaf dimensions. The function used to calculate functional diversity (dbFD) performs a Principal Coordinate Analysis on the traits and removes information redundancy. We also calculated species richness and total abundance (sum of the number of individuals per species).

Finally, we calculated the ratio between the values of each biodiversity metric in each plot assigned to an anthropogenic land use and its paired plot assigned to a natural habitat (hereafter called “relative functional richness”, “relative functional evenness”, “relative functional divergence”, “relative species richness” and “relative total abundance”) as follows:

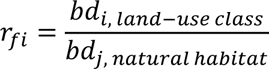

Where bd_i, anthropogenic land-use class_ is the biodiversity metric in anthropogenic vegetation plot *i* and bd_j, natural habitat_ is the biodiversity metric in the natural vegetation plot *j* paired to plot *i*. We also calculated the relative number of species shared between each plot *i* in anthropogenic land and the paired plot in natural habitats *j* compared to the total number of species in the plot in anthropogenic land (hereafter “relative natural species richness”). This metric is a relative value by definition and has the purpose of quantifying how much of the original composition contributes to the new species pool.

Relative values below one indicate that the absolute value is smaller in anthropogenic land use than in the natural habitat. Relative values above one indicate the opposite trend. To obtain aggregated values per bioregion, a median of all the relative values for each metric, land-use class and bioregion was calculated. The values per land-use class across bioregions were calculated as the weighted mean of the medians using the areas of bioregions as weights.

Given the assumptions made, we tested the sensitivity of the results by removing vegetation plots ranked 3 in terms of location uncertainty, imposing an upper threshold on the distance between PC scores of 0.01 (this threshold was selected, as it would remove most of data dispersion, Fig. S9.1), and excluding vegetation plots in anthropogenic land with a naturalness level of 2. The Wilcoxon signed-rank test for paired data with the Benjamini & Hochberg (hereafter “BH”) correction for multiple comparisons was used to determine the significance of the shift between the absolute values of biodiversity metrics in the paired plots.

## 3. Results

The final selection consisted of 73,074 vegetation plots (Fig. S11.1) and 7,185 vascular plant species, out of an estimated 20,000–25,000 European species (Bilz et al. 2011; POWO 2021) (see Fig. S8.2 and Fig. S9.1 for the spatial and density distribution of the plots after the pairing and for the ridge plot of the PC distance, respectively).

In most bioregions, we found lower relative functional richness and divergence (r_fi_ < 1) but higher relative functional evenness (r_fi_ > 1) in anthropogenic land than in the corresponding paired natural habitats (see Fig. 2, Tables S17-S19).

The difference in functional diversity between anthropogenic land-use classes and their paired natural counterparts was much more evident for functional richness than for functional evenness and divergence. On average, across all bioregions and land-use classes, functional richness and divergence were, respectively, 66% and 4% lower in anthropogenic areas compared to natural habitats. Functional evenness was 6% higher in anthropogenic areas than in natural ones.

The results of the sensitivity analysis did not significantly differ from the default settings (Figs. S12.1-S12.3).

### 3.1 Variation among land-use classes

Overall, *Cropland* and *Urban areas* were the land-use classes with the lowest relative functional richness compared to their paired natural vegetation (Fig. 2 and Fig. 3), especially in PA8 *Ural Mountains & West Eurasian Taiga Forests* and PA9 *Great Britain, Ireland & Faroe Islands*, where their values ranged between 0.08 and 0.1 (Table S18).

**Figure 3.**
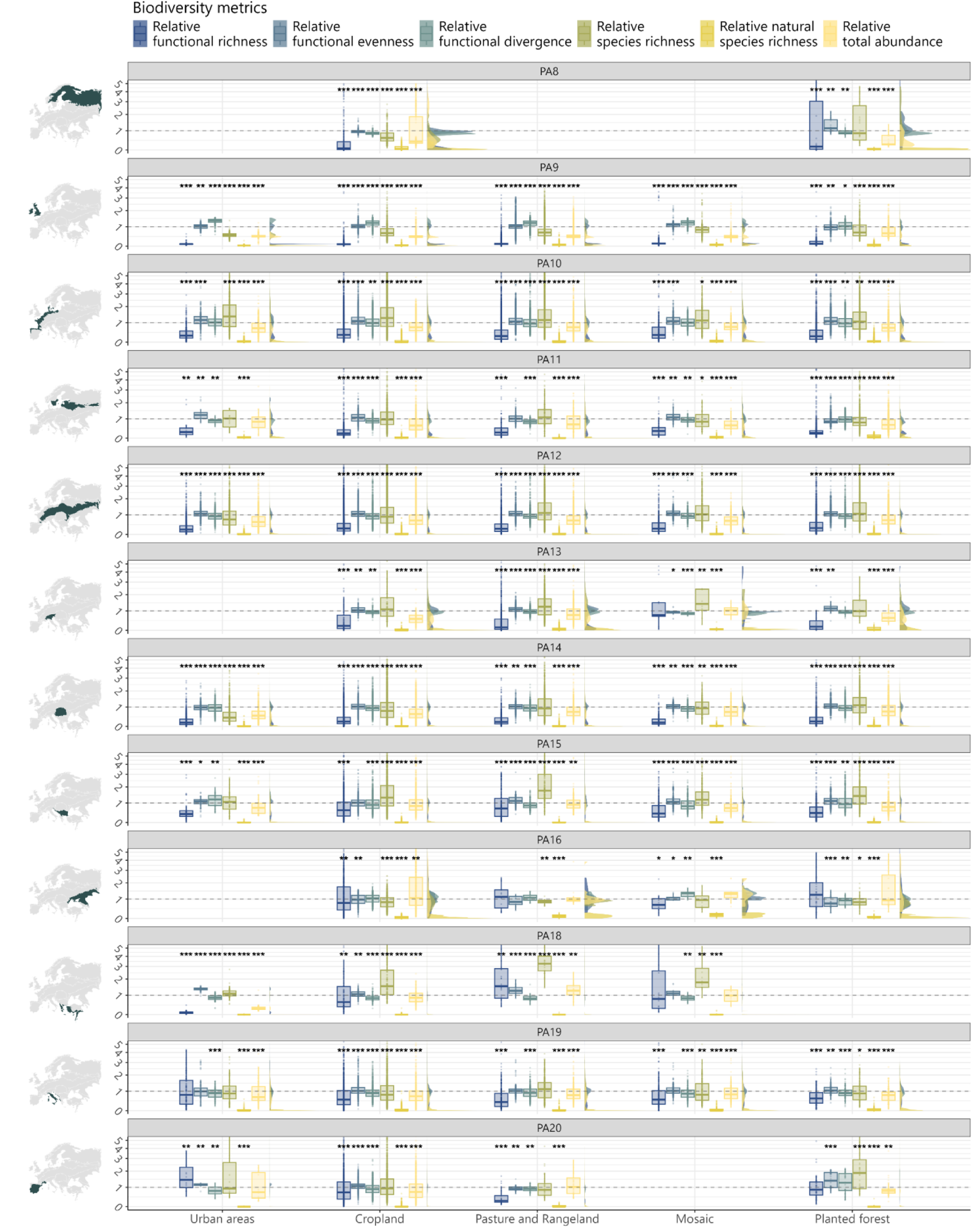
Relative functional and species diversity per bioregion and per land-use class, zoomed between 0 and 5 to improve readability. The dotted lines correspond to 1. Results of Wilcoxon test: ***: p-value ≤ 0.001, **: 0.001 < p-value ≤ 0.05, *: 0.05 < p-value ≤ 0.1, no *: p-value > 0.1.

Averaging the relative functional richness values per land-use class across bioregions (Table 2), we found the lowest weighted mean for *Cropland* and the highest one for *Pasture and rangeland*. *Cropland* and *Pasture and rangeland* were also the categories with the lowest minimum value and the highest maximum value, respectively. *Mosaic* showed the most homogeneous distribution of values of relative functional richness across bioregions (see Fig. 2 and relative weighted standard deviation in Table 2).

**Table 2.** Mean, minimum, maximum, and standard deviation of relative functional diversity values averaged per land-use class. The alphanumeric codes in brackets correspond to the bioregion where the minimum/maximum was found.

| Land-use class | Metric | Weighted mean | Minimum | Maximum | Relative weighted standard deviation |
| --- | --- | --- | --- | --- | --- |
| Urban areas | Relative functional richness | 0.38 | 0.10 (PA9) | 1.44 (PA20) | 1.02 |
|  | Relative functional evenness | 1.12 | 0.98 (PA14) | 1.36 (PA18) | 0.09 |
|  | Relative functional divergence | 1.01 | 0.81 (PA20) | 1.36 (PA9) | 0.18 |
| Cropland | Relative functional richness | 0.32 | 0.08 (PA8) | 0.79 (PA16) | 0.75 |
|  | Relative functional evenness | 1.02 | 0.95 (PA8) | 1.10 (PA10) | 0.05 |
|  | Relative functional divergence | 0.95 | 0.84 (PA18) | 1.21 (PA9) | 0.12 |
| Pasture and Rangeland | Relative functional richness | 0.46 | 0.11 (PA9) | 1.52 (PA18) | 0.93 |
|  | Relative functional evenness | 1.03 | 0.85 (PA16) | 1.25 (PA18) | 0.09 |
|  | Relative functional divergence | 0.97 | 0.82 (PA18) | 1.21 (PA9) | 0.13 |
| Mosaic | Relative functional richness | 0.39 | 0.13 (PA9) | 0.80 (PA18) | 0.55 |
|  | Relative functional evenness | 1.08 | 0.95 (PA13) | 1.11 (PA9) | 0.03 |
|  | Relative functional divergence | 1.02 | 0.83 (PA15) | 1.34 (PA16) | 0.17 |
| Planted forest | Relative functional richness | 0.37 | 0.12 (PA9) | 1.26 (PA16) | 0.87 |
|  | Relative functional evenness | 1.05 | 0.75 (PA16) | 1.38 (PA20) | 0.14 |
|  | Relative functional divergence | 0.95 | 0.88 (PA8) | 1.26 (PA20) | 0.10 |

*Cropland* had the lowest weighted mean also for relative functional divergence (together with *Planted forest*) and relative functional evenness. Relative functional divergence and relative functional evenness showed the highest weighted mean, respectively, in *Mosaic* and in *Urban areas.* The highest variation in terms of impact on functional evenness was found for *Planted forest* (the values are the most heterogeneous across bioregions, see Fig. 2 and weighted relative standard deviation in Table 2). We found the opposite for *Mosaic:* similar values of functional evenness in all bioregions and almost all above one, except PA13 *Alps & Po Basin Mixed Forests*.

### 3.2 Variation among bioregions

The variation of the response appeared to be more pronounced among bioregions than among land-use classes. PA9 *Great Britain, Ireland & Faroe Islands* showed the most homogeneous values of functional diversity and the lowest values of median relative functional richness for all land-use classes except *Cropland*, which had its minimum in *PA8 Ural Mountains & West Eurasian Taiga Forests* (Fig. 3). At the same time, PA9 depicted the highest values of relative functional divergence in *Urban areas, Cropland* and *Pasture and rangeland*. PA16 *Pontic Steppe Grasslands* had the largest number of land-use classes with relative functional richness above one (*Pasture and rangelands* and *Planted forest*) and did not show significant difference from natural habitat according to the Wilcoxon test. PA18 *Aegean Sea & East Mediterranean Mixed Forests*, PA19 *Adriatic Sea & Central Mediterranean Mixed Forests* and PA20 *Balearic Sea & West Mediterranean Mixed Forests*, all in southern Europe, had higher values of relative functional richness compared to the more northern bioregions (Fig. 2), even above one in two cases, resulting from a lower absolute value of functional richness for the two natural vegetation classes occurring there, *Natural forest* and *Natural shrubland* (Fig. S13.1).

### 3.3 Comparison with other biodiversity metrics

The impact of anthropogenic land use on relative species richness did not yield the same results as those of relative functional richness (Fig. 3), and their relationship followed a logarithmic pattern (see Figs. S14.1 and S14.2). In 25 combinations of land-use classes and bioregions, the medians of both relative species richness and relative functional richness were below one, and, in most cases, the decrease in functional richness appeared more pronounced. In 23 other combinations, especially for the land-use class *Pasture and rangelands*, the median relative species richness was around one or slightly above one, whereas the median relative functional richness was below one. Conversely, in three combinations distributed in PA16 *Pontic Steppe Grasslands* and in PA20 *Balearic Sea & West Mediterranean Mixed Forests*, the opposite occurred. *Pasture and rangeland* in PA18 *Aegean Sea & East Mediterranean Mixed Forests* was the only case where both metrics were above one.

The relative natural species richness (Fig. 3) was very close to zero for all combinations of anthropogenic land-use classes and bioregions and the relative total abundance had a median value smaller than one in most combinations of land-use classes and bioregions (Fig. 3), with a few exceptions in southern Europe.

## 4. Discussion

We found evidence that anthropogenic land use leads to a reduction in functional richness and divergence while simultaneously promoting an increase in functional evenness. This trend was consistent across most of the bioregions we examined, with a particularly pronounced effect observed for functional richness. Our results confirm those of Scherer *et al*. (2020) at a smaller spatial extent in Europe but diverge from the findings of Flynn *et al*. (2008), who reported no clear pattern of diversity change due to anthropogenic land use for plants in the Americas. The disparity in results may be attributed to differences in the metric used for assessing functional diversity. Flynn *et al*. (2008), along with De Souza *et al*. (2014), which is mostly based on data from Flynn *et al*. (2008), calculated a single functional diversity metric, which was quantified based on a dendrogram and relied on only eight functional traits, including four categorical traits, found to decrease the quality of the measure (Maire et al. 2015).

Regarding the response of each functional diversity metric, the observed decrease in functional richness aligns with previous studies (Pakeman 2011; Janeček et al. 2013; Carmona et al. 2020): human disturbance occurring with anthropogenization of land tends to diminish functional richness. Concerning functional divergence, previous studies have reported mixed responses to land use (Bonilla-Valencia et al. 2022). The decrease recorded in our study suggests that anthropogenic land use reduces the degree of niche differentiation among species within communities and increases the trait similarity of dominant species (Mason et al. 2005; Mouchet et al. 2010). Conversely, human disturbance appeared to promote functional evenness. Functional evenness is higher when species and their abundance are more regularly distributed along functional trait gradients (Mouchet et al. 2010; Pakeman 2011). When coupled with a reduction of traits’ multidimensional space (reduced functional richness) and with a lower degree of niche differentiation (reduced functional divergence), the response of functional evenness suggests that the dominant species exhibit a higher level of similarity in anthropogenic land than in natural habitats (Mouchet et al. 2010; Pakeman 2011). These findings may indicate that species populating anthropogenic areas are more functionally homogeneous than those in natural habitats. Previous studies have reported disruption of interactions among specialized partners following the removal of natural and semi-natural habitats, resulting in extensive community restructuring towards a less diverse community dominated by generalist and widespread species (Newbold 2018; Le Provost et al. 2021). These effects are likely linked to the extensive change in species composition observed when comparing anthropogenic land use with natural habitats (Fig. 3). For example, plots in anthropogenic land in PA16 *Pontic Steppe Grasslands* had higher relative natural species richness and functional richness than other bioregions, suggesting that species turnover plays a big role in determining functional richness under disturbance.

The logarithmic relationship between species richness and functional richness has been observed in prior studies (Villéger et al. 2008; Biswas and Mallik 2011; Aros-Mualin et al. 2021; Boonman et al. 2021), likely accounting for the occasionally divergent behavior observed in these metrics. Notably, natural habitats tended to exhibit higher functional richness than anthropogenic land use at equivalent levels of species richness (Fig. S14.1). Moreover, as species richness increased, the corresponding rise in functional richness did not occur at the same rate. Conversely, lower species richness was associated with more pronounced variations in functional richness. This underscores the distinct information conveyed by these indices and emphasizes how functional diversity can detect adverse changes even when species richness remains unchanged or seemingly improves.

Regarding species richness alone, we acknowledge that, in contrast with our findings, many studies with site-specific comparisons focusing on populations have found evidence of decline in all land-use classes (Newbold et al. 2015). In addition, there are inherent uncertainties in our study due to the quality of the datasets, which suggests pairing refinement should be prioritized once additional data becomes available (see Limitations section). Compared to previous studies, we implemented a different and more systematic approach, not based on meta-analyses but on harmonized datasets and coupling anthropogenic land with the potential natural vegetation occurring there and not the natural vegetation nearby, which might differ from the original natural vegetation. The differences may emerge from these elements. Furthermore, a global review of land-use effects on plant species richness found a positive trend in managed grasslands compared to natural habitats (Gerstner et al. 2014). Additionally, recent research indicates that global median species richness may either increase or remain stable when comparing different types of natural forests (namely, natural deciduous broadleaf forest, evergreen needleleaf forests, and mixed forests, which are the same natural forests considered in our study) to minor or major human modification (Deng et al. 2024). Furthermore, Flynn *et al*. (2009) did not find a significant difference between plant species richness in anthropogenic and natural land.

Variability in response among bioregions, compared to land-use classes, may stem from differences in environmental conditions and management practices. Environmental factors can act as filters on trait composition within plant communities, influencing their functional diversity (Bruelheide et al. 2018; Wallis et al. 2021; Cheng et al. 2022; Joswig et al. 2022). While PCA was utilized to minimize this influence when pairing vegetation plots, distinct levels of functional variation may still be exhibited across bioregions due to their specific natural vegetation and environment. For instance, Mediterranean bioregions (PA18, PA19, PA20) exhibited lower absolute functional richness compared to northerly ones (Fig. S13.1). Certain forest types, such as mixed forests (mostly spread in central Europe) and coniferous forests (occurring at high latitudes or high elevations, e.g. in PA8 or PA13, which we recorded having the highest absolute values of functional richness, Fig. 13.1), can exhibit larger trait hypervolumes than Mediterranean woodlands or temperate grasslands (Echeverría-Londoño et al. 2018), corroborating our findings. Furthermore, the relationship between trait hypervolume and latitude has been recorded to differ from that of species richness, indicating distinct ecological dynamics (Lamanna et al. 2014).

Additionally, management practices and intensity within land-use classes vary across bioregions and likely modulate their impacts on functional diversity (Laliberté et al. 2010; Janeček et al. 2013; Van Meerbeek et al. 2014). While our study could not incorporate management intensity due to data limitations, future research should consider this aspect for a more comprehensive assessment (Dullinger et al. 2021).

Our study’s findings on how human land use affects plant functional diversity have significant implications for future global change. Predictive models suggest that trait evolution aids resilience (Guerin et al. 2014), while the threat of invasive alien species, amplified by global trade, is influenced by the functional diversity and abundance of native species (Tordoni et al. 2020; Kaushik et al. 2022; Díaz et al. 2023). For instance, the changes we observed, like decreased functional richness and divergence alongside increased functional evenness due to human pressures, may alter the native communities’ ability to resist invasive species. Shifts in functional diversity may also heighten plant communities’ response to nitrogen deposition, particularly relevant in Europe, where it fosters nitrogen-demanding plants (Staude et al. 2020). Considering these observed patterns, we anticipate ongoing alterations in plant functional diversity dynamics under global changes such as land use change.

### Limitations

To evaluate the effects of human land use on vascular plant diversity comprehensively, we had to face a few challenges. Firstly, data coverage varied across bioregions and land-use classes, particularly in southern Europe and the Mediterranean region, where the number of natural vegetation plots available for analysis was limited by data availability and by the long history of land management (Martín-Forés 2017; Sabatini et al. 2018; Ellis et al. 2021). Secondly, the absence of a consistent spatial data source for land-use classification led us to combine multiple maps, introducing uncertainty. The main challenge lays in identifying a reliable source capable of distinguishing between natural and managed forests at a meaningful spatiotemporal resolution, which remains difficult despite advancements in remote sensing (Hirschmugl et al. 2017). Thirdly, the matching between the vegetation plots and the land-use and land-cover classes introduced uncertainty, mainly because of three factors: (i) the location uncertainty of the vegetation plots (see Fig. S15.2), which we addressed by excluding plots with high location uncertainty and ranking the others based on the homogeneity of land use/cover within the uncertainty radius (see sec. 2.1); (ii) the intrinsic uncertainty of the land-use and land-cover maps; and (iii) the mismatch between the resolution of the land-use map and the size of the vegetation plots (53 m^2^ on average but with much higher or not available values for part of the plots (see Fig. S15.1), which could lead to incorrect land-use assignments.

### Outlook

Our analysis stands out for its comprehensive consideration of a high number of traits, surpassing previous studies. To increase even more the trait representativeness, enhancing belowground trait coverage is advisable (Carmona, Bueno, et al. 2021). Exploring solutions for trait distribution gaps or scaling up with methods like model predictions or remote sensing is worth investigating (Schneider et al. 2017; Boonman et al. 2020; Hauser, Timmermans, et al. 2021; Hauser, Féret, et al. 2021).

An improvement in the ecological relevance of the assessment and an in-depth comprehension of the effect of land-use change from natural to human-modified could be achieved by considering not only plants’ traits but also traits of other taxonomic groups. This advancement would make it possible to trace the effect along the trophic chain and on species interactions, given the essential interplay between them (Haddad et al. 2009; Rigal et al. 2023; Windsor et al. 2023).

## Conclusion

Our study offers valuable insights into the multifaceted relationship between anthropogenic land modification and biodiversity and highlights the significance of incorporating diverse metrics, notably functional diversity, which yields unique and complementary insights beyond traditional measures. We integrate information across different databases, enabling data interoperability and merging various sources of information at a continental scale. Through our regionalized approach and novel methodology, we enhanced our understanding of this intricate relationship and introduced a fresh perspective on connecting localized studies with broader regional implications. Such efforts are crucial in addressing the urgent challenge of halting biodiversity loss.

## Code and data availability

The code and the data on which the analysis is based are available for the review process on Dryad at: https://datadryad.org/stash/share/50nM9Rbd-qMSoiREeLir9CekoiFLG39bt3V5PiUdj4s

