## Supplementary material for "Land-use impacts on plant functional diversity throughout Europe": SI

Francesca Rosa, ETH Zürich

Institute of Environmental Engineering, chair of Ecological Systems Design

HIF D 11, Laura-Hezner-Weg 7, 8093 Zürich, Switzerland

+41 44 633 99 49

### S1 Flowchart of the methodology

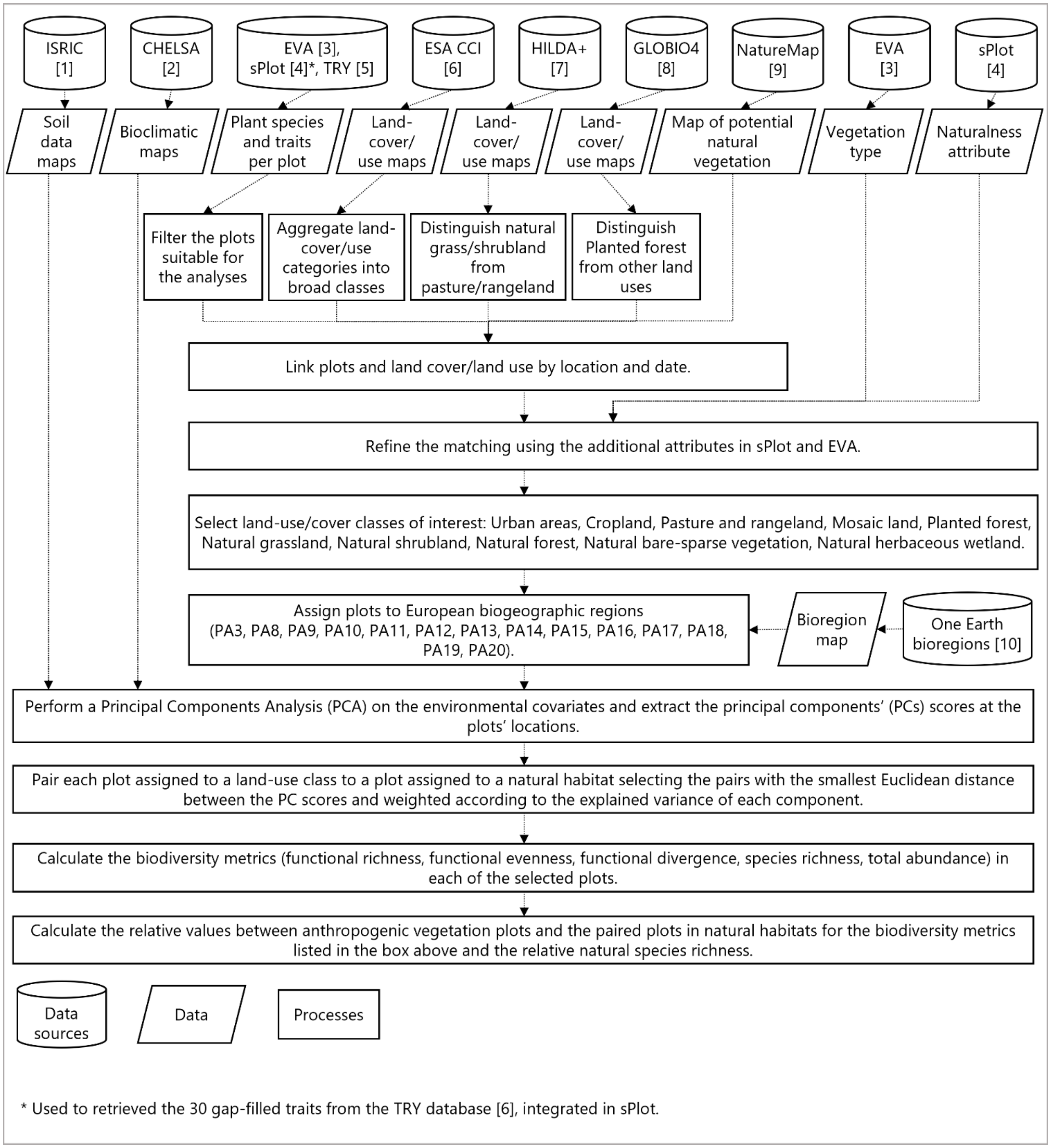


Figure S1.1 Flowchart of methodology applied in this study to quantify relative functional and species diversity.

1. (ISRIC; Hengl et al. 2017)
2. (Karger et al. 2017; Karger et al. 2018)
3. (Chytrý et al. 2016)
4. (Bruelheide et al. 2019)
5. (Kattge et al. 2020)
6. (ESA Land Cover CCI project team and Defourny 2019)
7. (Winkler et al. 2021)
8. (Schipper et al. 2019; PBL Netherlands Environmental Assessment Agency 2023)
9. (Hengl et al. 2020)
10. (One Earth 2020)

#### S2 List of filtering criteria

Filtering criteria on vegetation plots.

All the following criteria had to be fulfilled for a plot to be kept in the selection.

- Availability of
    - Coordinates
    - Date of recording
    - Information on abundance (expressed as cover percentage)
    - Information on level of naturalness (needed for the land use matching, see Sec. 2.3 and 2.4)
- Attributes in the following range:
    - Traits information for at least 3 species per plot
    - Proportion of species with traits (PST) $\geq 0.5$ and trait coverage (TC) $\geq 0.8$
    - Range of plot size between 1 $m^2$ and 500 $m^2$
    - Location uncertainty < 150 m or not available or location uncertainty < $10^4$ m with at least 80% homogenous land use within the uncertainty radius (and land use class equal to the one assigned by the land use matching, see Sec. 2.4).

Filtering criteria on species:

- Only vascular plants
- Only species with traits

#### S3 Bioregions

Table S3.1 List of European bioregions

| Code | Name | Code | Name |
|---|---|---|---|
| **PA3** | Scandinavian Birch & Coastal Conifer Forests | PA14 | Carpathian Mountain & Plains Mixed Forests |
| **PA8** | Ural Mountains & West Eurasian Taiga Forests | PA15 | Dinaric Mountains & Balkan Mixed Forests |
| **PA9** | Great Britain, Ireland & Faroe Islands | PA16 | Pontic Steppe Grasslands |
| **PA10** | West European Coastal Mixed Forests | PA17 | Black Sea, Caucasus-Anatolian Mixed Forests & Steppe |
| **PA11** | Baltic Sea & Sarmatic Mixed Forests | PA18 | Aegean Sea & East Mediterranean Mixed Forests |
| **PA12** | European Interior Mixed Forests | PA19 | Adriatic Sea & Central Mediterranean Mixed Forests |
| **PA13** | Alps & Po Basin Mixed Forests | PA20 | Balearic Sea & West Mediterranean Mixed Forests |

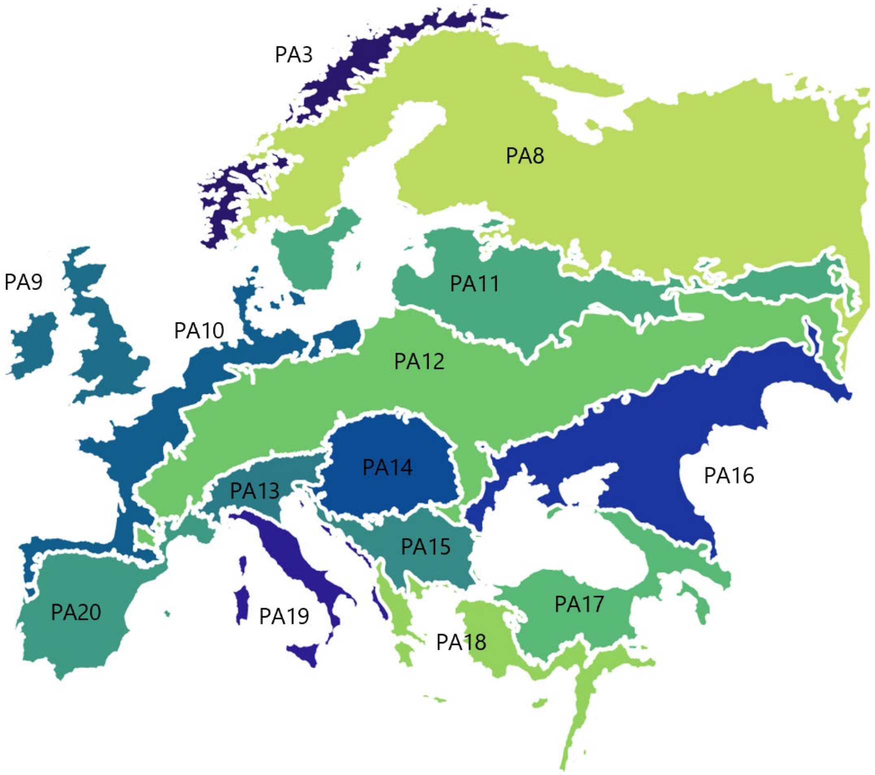


Figure S3.1 Map of European bioregions (One Earth 2020).

#### S4 Aggregation of land use/cover of ESA CCI

Table S4.1 Matching between the land-use/cover classes of ESA CCI and the land-use/cover classes defined in the process of aggregating multiple categories and reaching the land-use/cover classification suitable for the study (see Sec. 2.3).

| ESA CCI code | ESA CCI label | Corresponding broad class |
|---|---|---|
| **10** | Cropland rainfed | Cropland |
| **11** | Cropland rainfed, Herbaceous cover | Cropland |
| **12** | Cropland rainfed, Tree or shrub cover | Cropland |
| **20** | Cropland irrigated or post-flooding | Cropland |
| **30** | Mosaic cropland (>50%) / natural vegetation (tree shrub herbaceous cover) (<50%) | Mosaic_managed |
| **40** | Mosaic natural vegetation (tree shrub herbaceous cover) (>50%) / cropland (<50%) | Mosaic_managed |
| **50** | Tree cover broadleaved evergreen closed to open (>15%) | Forest |
| **60** | Tree cover broadleaved deciduous closed to open (>15%) | Forest |
| **61** | Tree cover broadleaved deciduous closed to open (>15%), Tree cover broadleaved deciduous closed (>40%) | Forest |
| **62** | Tree cover broadleaved deciduous closed to open (>15%), Tree cover broadleaved deciduous open (15-40%) | Forest |
| **70** | Tree cover needleleaved evergreen closed to open (>15%) | Forest |
| **71** | Tree cover needleleaved evergreen closed to open (>15%), Tree cover needleleaved evergreen closed (>40%) | Forest |
| **72** | Tree cover needleleaved evergreen closed to open (>15%), Tree cover needleleaved evergreen open (15-40%) | Forest |
| **80** | Tree cover needleleaved deciduous closed to open (>15%) | Forest |
| **81** | Tree cover needleleaved deciduous closed to open (>15%), Tree cover needleleaved deciduous closed (>40%) | Forest |
| **82** | Tree cover needleleaved deciduous closed to open (>15%), Tree cover needleleaved deciduous open (15-40%) | Forest |
| **90** | Tree cover mixed leaf type (broadleaved and needleleaved) | Forest |
| **100** | Mosaic tree and shrub (>50%) / herbaceous cover (<50%) | Mosaic_natural |
| **110** | Mosaic herbaceous cover (>50%) / tree and shrub (<50%) | Mosaic_natural |
| **120** | Shrubland | Shrubland |
| **121** | Shrubland, Shrubland evergreen | Shrubland |
| **122** | Shrubland, Shrubland deciduous | Shrubland |
| **130** | Grassland | Grassland |
| **140** | Lichens and mosses | Lichens_and_mosses |
| **150** | Sparse vegetation (tree shrub herbaceous cover) (<15%) | Sparse_vegetation |

| | | |
|---|---|---|
| **151** | Sparse vegetation, Sparse tree (<15%) | Sparse_vegetation |
| **152** | Sparse vegetation, Sparse shrub (<15%) | Sparse_vegetation |
| **153** | Sparse vegetation, Sparse herbaceous cover (<15%) | Sparse_vegetation |
| **160** | Tree cover flooded fresh or brakish water | Tree_cover_flooded_land |
| **170** | Tree cover flooded saline water | Tree_cover_flooded_land |
| **180** | Shrub or herbaceous cover flooded fresh/saline/brakish water | Herbaceous_wetland |
| **190** | Urban areas | Urban |
| **200** | Bare areas | Bare_area |
| **201** | Consolidated bare areas | Bare_area |
| **202** | Unconsolidated bare areas | Bare_area |
| **210** | Water bodies | Water_bodies |
| **220** | Permanent snow and ice | Snow_and_ice |

#### S5 Aggregation of land cover of NatureMap (Potential natural vegetation)

Table S5.1 Matching between the land cover classes of NatureMap and the land use/cover classes defined in the process of aggregating multiple categories and reaching the land cover classification of potential natural vegetation suitable for the study (see Sec. 2.3).

| NatureMap code | NatureMap label | Corresponding broad class |
|---|---|---|
| **111** | closed forest, evergreen needleleaf | Forest_natural |
| **113** | closed forest, deciduous needleleaf | Forest_natural |
| **112** | closed forest, evergreen broadleaf | Forest_natural |
| **114** | closed forest, deciduous broadleaf | Forest_natural |
| **115** | closed forest, mixed | Forest_natural |
| **116** | closed forest, unknown | Forest_natural |
| **121** | open forest, evergreen needleleaf | Forest_natural |
| **123** | open forest, deciduous needleleaf | Forest_natural |
| **122** | open forest, evergreen broadleaf | Forest_natural |
| **124** | open forest, deciduous broadleaf | Forest_natural |
| **125** | open forest, mixed | Forest_natural |
| **126** | open forest, unknown | Forest_natural |
| **21** | sub-polar or polar barren-lichen-moss, grassland | Lichens_and_mosses |
| **20** | shrubs | Shrubland_natural |
| **30** | herbaceous vegetation | Grassland_natural |
| **90** | herbaceous wetland | Herbaceous_wetland |
| **100** | moss and lichen | Lichens_and_mosses |
| **60** | bare/sparse vegetation | Bare_Sparse_vegetation |
| **40** | cropland | Cropland |
| **50** | urban/built-up | Urban |
| **70** | snow and ice | Snow_and_ice |
| **80** | permanent water bodies | Water_bodies |
| **200** | open sea | Sea |

#### S6 Final land-use classification

Table S6.1 Overview of the matching between the various land use/cover spatial data and of the sPlot attributes used to derive the final land-use/cover classification assigned to the vegetation plots.

| Land cover/use assigned to the plots | ESA CCI name and code | HILDA+ | GLOBIO4 | sPlot veg. Type ** | sPlot Natural-ness ** | Natural habitats (N) or Human land use (H) |
|---|---|---|---|---|---|---|
| Natural forest | All tree cover types (50-90) | - | - | Forest | 1 | N |
| Natural grassland | Grassland (130) | Natural grass/ shrubland | - | Grassland | 1 | N |
| | Mosaic herbaceous cover (>50%) / tree and shrub (<50%) (110)* | - | - | - | | |
| Natural shrubland | All types of shrubland (120-122) | Natural grass/ shrubland | - | Shrubland | 1 | N |
| | Mosaic tree and shrub (>50%) / herbaceous cover (<50%) (100)* | - | - | - | | |
| Natural herbaceous wetland | Shrub or herbaceous cover flooded fresh/saline/brakish water (180) | - | - | Wetland | 1 | N |
| Natural bare and sparse vegetation | Sparse vegetation | - | - | Sparse vegetation | 1 | N |
| | All bare area types (200-202) | - | - | | | |
| Planted forest | - | - | Forestry | - | 2 or 3 | H |
| Pasture and rangeland | Grassland (130)<br>All types of shrubland (120-122) | Pasture/ Rangeland | - | - | 2 or 3 | H |
| Cropland | All cropland types (10-20) | - | - | - | 2 or 3 | H |
| Urban areas | Urban areas (190) | - | - | - | 2 or 3 | H |
| Mosaic | Mosaic cropland (>50%) / natural vegetation (tree shrub herbaceous cover) (<50%) (30)<br>Mosaic natural vegetation (tree shrub herbaceous cover) (>50%) / cropland (<50%) (40) | - | - | - | 2 or 3 | H |

* In the Mediterranean region, and especially in the Iberian Peninsula, the ESA CCI map does not assign any plot to the class *Shrubland*, to which instead the NatureMap assigns a large area. To prevent losing all the plots that would be paired to the control *Natural shrubland* but for which a control would not be found the class *Mosaic tree and shrub (>50%) / herbaceous cover (<50%)* was considered as Natural shrubland, since many plots are assigned to that class by the ESA CCI map. The same approach was applied to *Mosaic herbaceous cover (>50%) / tree and shrub (<50%)* for *Natural grassland.*
** This is primary information contributed by the owner of every dataset in sPlot. The guidelines used by the data contributors to sPlot to give these definitions are available on the sPlot webpage (sPlot 2024a). Naturalness levels: 1 = natural, 2 = semi-natural, 3 = anthropogenic.

### S7 Environmental covariates and PCA

Table S7.1 List of bioclimatic variables and soil properties used as input for the PCA.

| **Bioclimatic variables (CHELSA V2.1) – Mean** | 13. bio13: Precipitation of Wettest Month |
|---|---|
| 1. bio01: Annual Mean Temperature | 14. bio14: Precipitation of Driest Month |
| 2. bio02: Mean Diurnal Range | 15. bio15: Precipitation Seasonality |
| 3. bio03: Isothermality | 16. bio16: Precipitation of Wettest Quarter |
| 4. bio04: Temperature Seasonality | 17. bio17: Precipitation of Driest Quarter |
| 5. bio05: Max Temperature of Warmest Month | 18. bio18: Precipitation of Warmest Quarter |
| 6. bio06: Min Temperature of Coldest Month | 19. bio19: Precipitation of Coldest Quarter |
| 7. bio07: Temperature Annual Range | **Soil Variables** (SoilGrids) – Mean |
| 8. bio08: Mean Temperature of Wettest Quarter | 20. CLYPPT: Clay mass fraction in % |
| 9. bio09: Mean Temperature of Driest Quarter | 21. PHIHOX: Soil pH x 10 in H20 |
| 10. bio10: Mean Temperature of Warmest Quarter | 22. SLTPPT: Silt mass fraction in % |
| 11. bio11: Mean Temperature of Coldest Quarter | 23. SNDPPT: Sand mass fraction in % |
| 12. bio12: Annual Precipitation | |

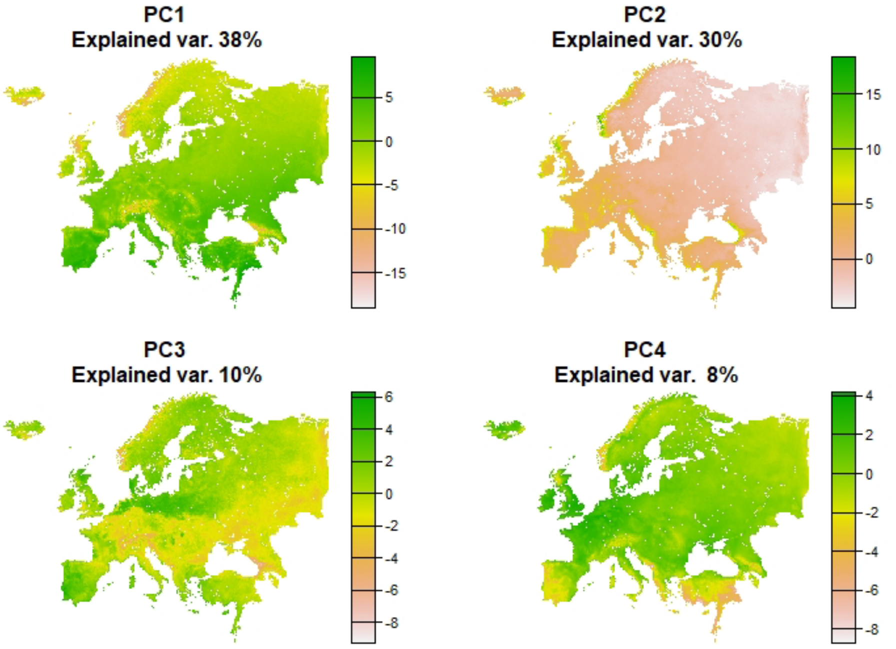


Figure S7.1 Spatial representation of the first four PCs that explain most of the variance.

#### S8 Pairing of vegetation plots

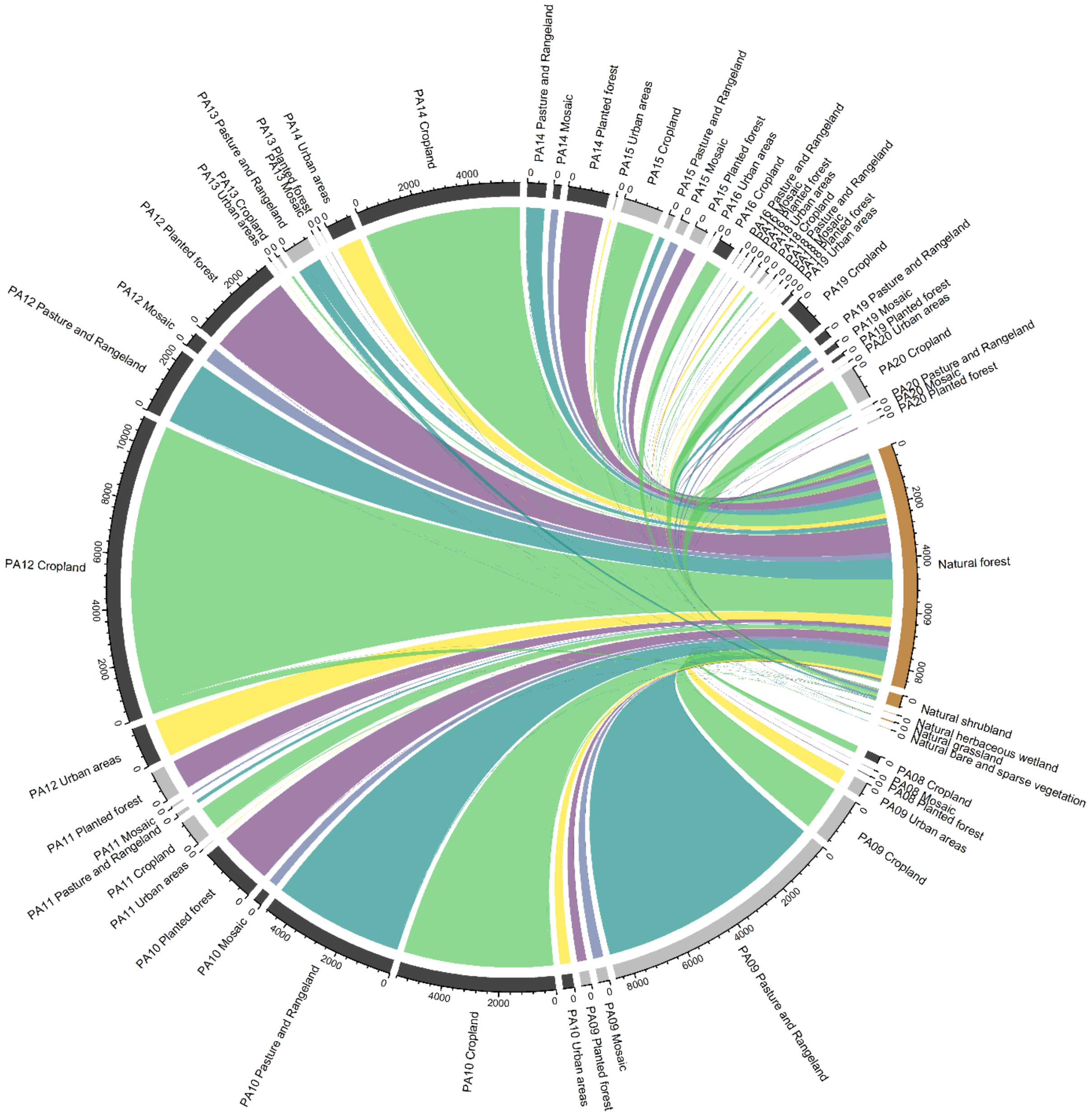


Fig. S8.1 Chord diagram (Gu 2014) displaying the pairing between plots in human-modified land (grouped by bioregion and land-use class) and plots in natural habitats (grouped by land cover). The number of plots in natural land is lower than in human-modified land classes because, as mentioned in the text, multiple plots in human-modified land could be paired with the same plot in natural habitat, having the minimum PC score distance.

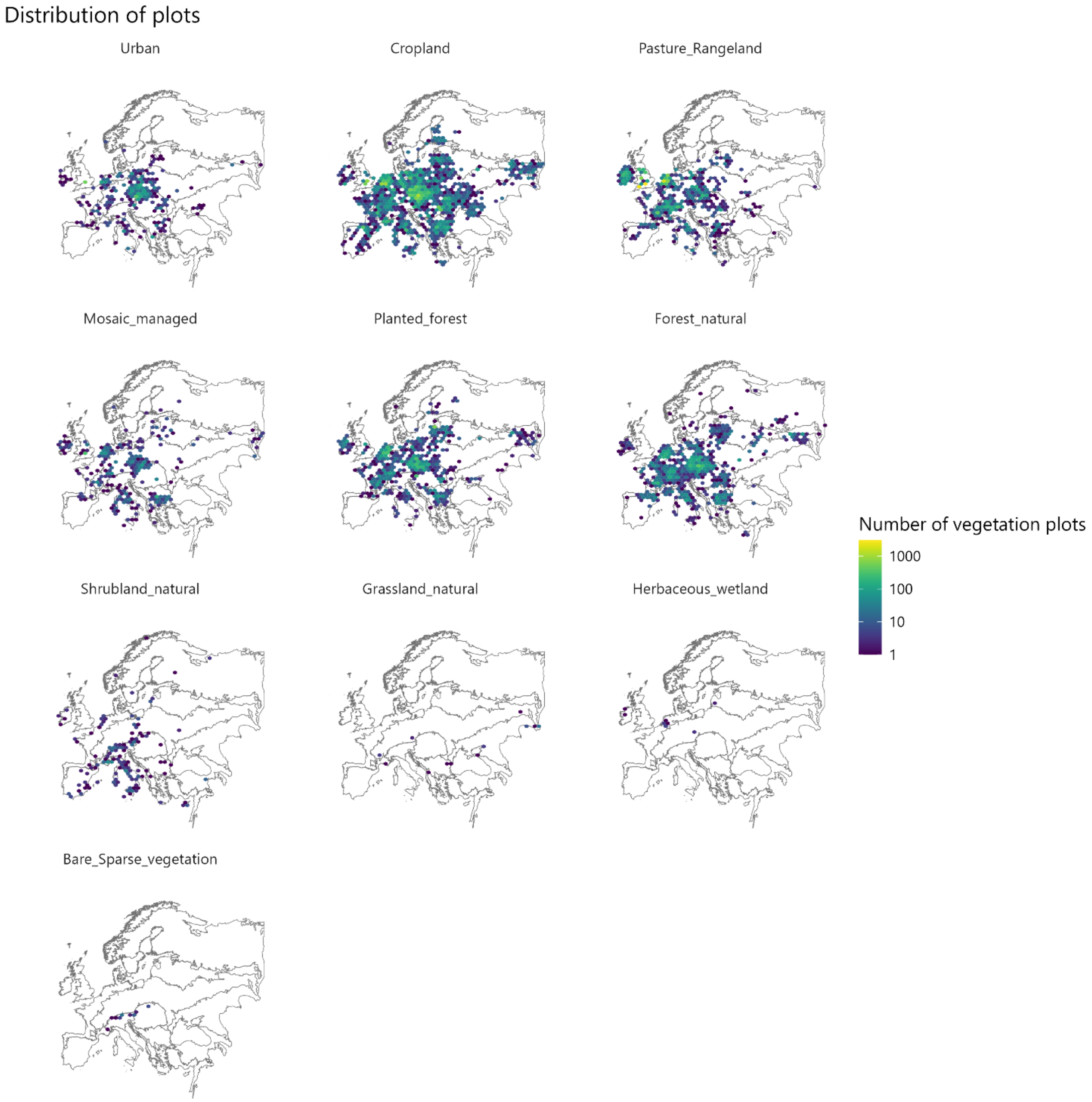


Figure S8.2 Spatial distribution and density of the plots remaining after the pairing process and considered in this study for the calculation of functional diversity.

#### S9 Distance of PC scores

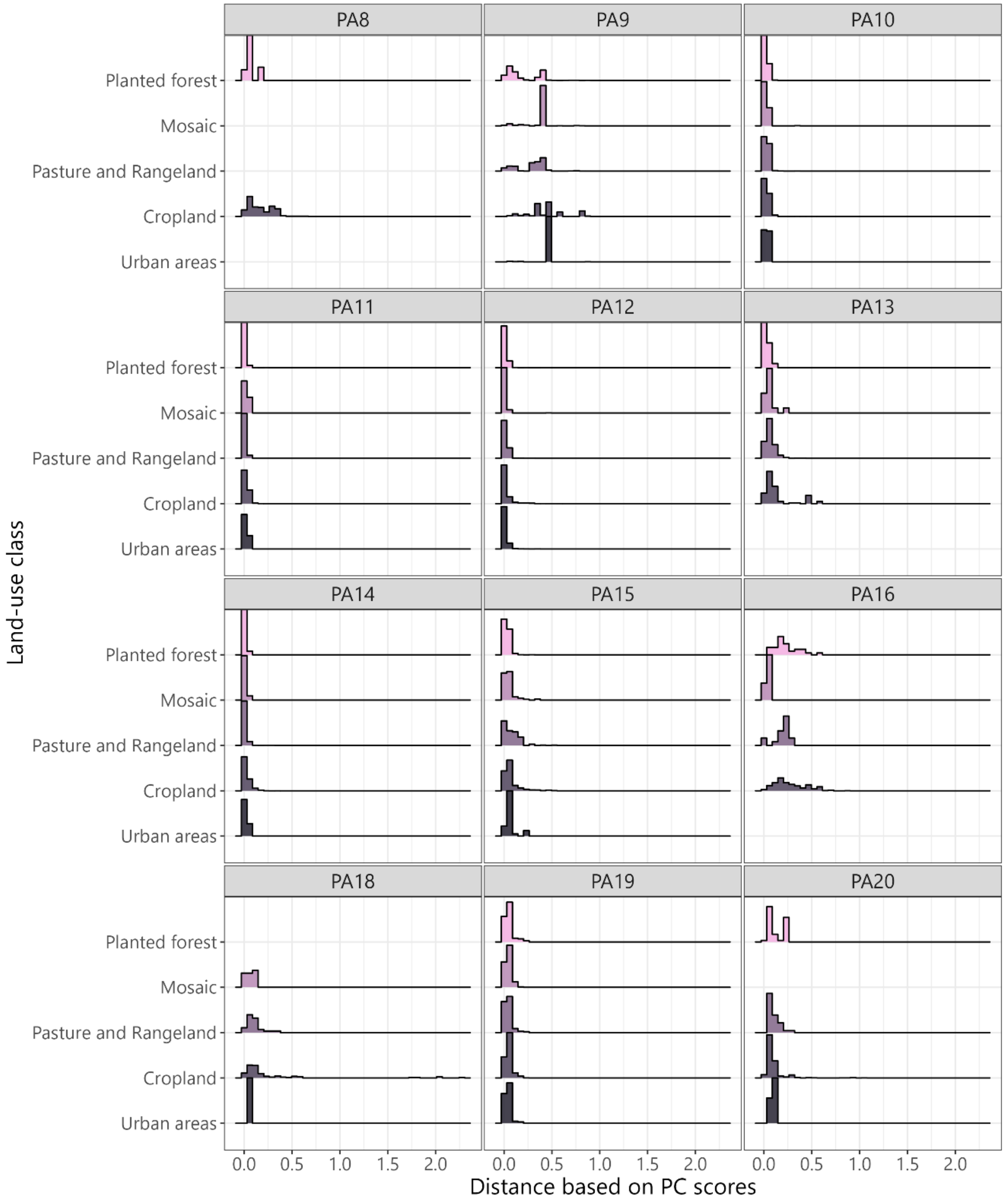


Figure S9.1 Ridge plot displaying the distribution of distances between the PC scores of plots in human-modified land and natural habitats after the pairing.

#### S10 Traits

Table S10.1 List of plant species traits considered in this study, also available in sPlot documentation (sPlot 2024b), with categorization.

| trait | code | full name | unit | Category based on (Falster et al. 2021) | Category based on (Kattge et al. 2011) |
|---|---|---|---|---|---|
| SLA | 11 | Leaf area per leaf dry mass (specific leaf area, SLA) | - | leaf allocation | morphology |
| LDMC | 47 | Leaf dry mass per fresh mass (leaf dry matter content, LDMC) | [g/g] | leaf allocation | anatomical |
| LeafWaterCont | 3120 | Leaf water content per dry mass (not saturated) | [g/g] | leaf allocation | anatomical |
| LeafThickness | 46 | Leaf thickness | [mm] | leaf morphology | morphology |
| LeafNperArea | 50 | Leaf nitrogen (N) content per leaf area | [g/m2] | leaf morphology | morphology |
| LeafDryMass.single | 55 | Leaf dry mass (single leaf) | [mg] | leaf morphology | morphology |
| LeafLength | 144 | Leaf length | [mm] | leaf morphology | morphology |
| LeafWidth | 145 | Leaf width | [cm] | leaf morphology | morphology |
| Leaffreshmass | 163 | Leaf fresh mass | [g] | leaf morphology | morphology |
| LeafArea.leaf.undef | 3112 | Leaf area (in case of compound leaves: leaf, undefined if petiole in- or excluded) | [$mm^2$] | leaf morphology | morphology |
| LeafC.perdrymass | 13 | Leaf carbon (C) content per leaf dry mass | [mg/g] | leaf nutrient | biochemical |
| LeafN | 14 | Leaf nitrogen (N) content per leaf dry mass | [mg/g] | leaf nutrient | biochemical |
| LeafP | 15 | Leaf phosphorus (P) content per leaf dry mass | [mg/g] | leaf nutrient | biochemical |
| LeafCN.ratio | 146 | Leaf carbon/nitrogen (C/N) ratio | - | leaf nutrient | biochemical |
| Leaf.delta.15N | 78 | Leaf nitrogen (N) isotope signature | [per] | leaf physiology | physiology |
| Seed.num.rep.unit | 138 | Seed number per reproduction unit | - | reproductive allocation | anatomical |
| SeedGerminationRate | 95 | Seed germination rate (germination efficiency) | [%] | reproductive life history | phenology |
| Chromosome.n | 223 | Species genotype: chromosome number | - | reproductive life history | phenology |
| Chromosome.cDNAcont | 224 | Species genotype: chromosome cDNA content | [pg] | reproductive life history | phenology |
| SeedMass | 26 | Seed dry mass | [mg] | reproductive morphology | morphology |
| Seed.length | 27 | Seed length | [mm] | reproductive morphology | morphology |
| Disp.unit.leng | 237 | Dispersal unit length [mm] | [mm] | reproductive morphology | morphology |
| SpecificRootLength | 1080 | Root length per root dry mass (specific root length, SRL) | [cm/g] | root allocation | anatomical |
| RootingDepth | 6 | Rooting depth | [m] | root morphology | morphology |
| StemDiam | 21 | Stem diameter | [m] | stem allocation | anatomical |
| StemDens | 4 | Stem specific density or wood density | [g/cm3] | stem morphology | morphology |
| PlantHeight | 18 | Plant height | [m] | stem morphology | morphology |
| Stem.cond.dens | 169 | Stem conduit density (vessels and tracheids) | [$mm^{-2}$] | stem morphology | morphology |
| StemConduitDiameter | 281 | Stem conduit diameter (vessels and tracheids) | [μm] | stem morphology | morphology |
| Wood.vessel.length | 282 | Wood vessel element length or stem conduit element length | [μm] | stem morphology | morphology |
| WoodFiberLength | 289 | Wood fiber lengths | [μm] | stem morphology | morphology |

#### S11 Heatmaps of the results with the number of plots per bioregion and land-use class

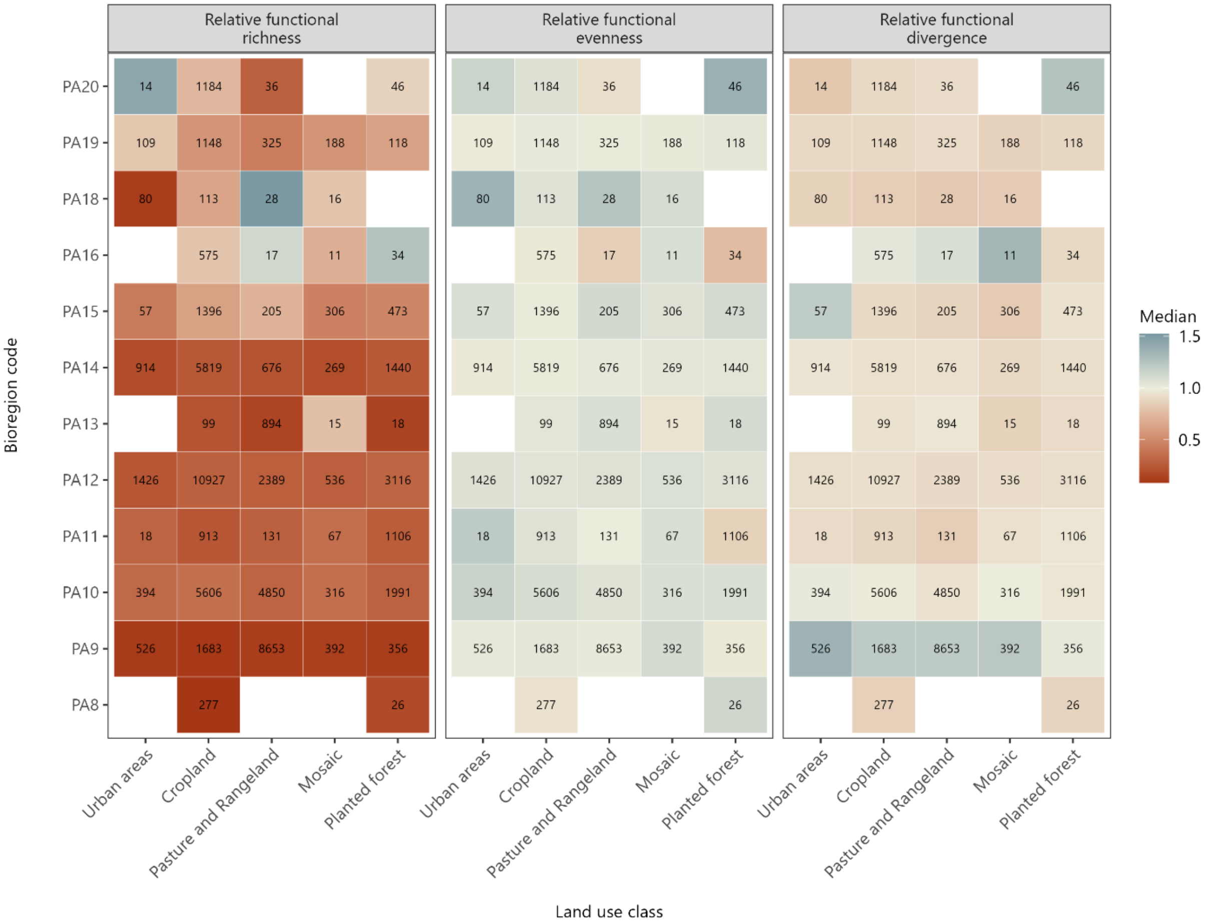


Figure S11.1 Relative functional richness, evenness and divergence per bioregion and human-modified land-use class with the same legend. The numbers in the cells correspond to the number of pairings. As described in the text, multiple plots in human-modified land may have been assigned to the same plot in natural habitat. Thus, the number of pairings reflects the number of unique plots in land used by humans but not the number of unique plots in natural habitat.

### S12 Sensitivity analysis

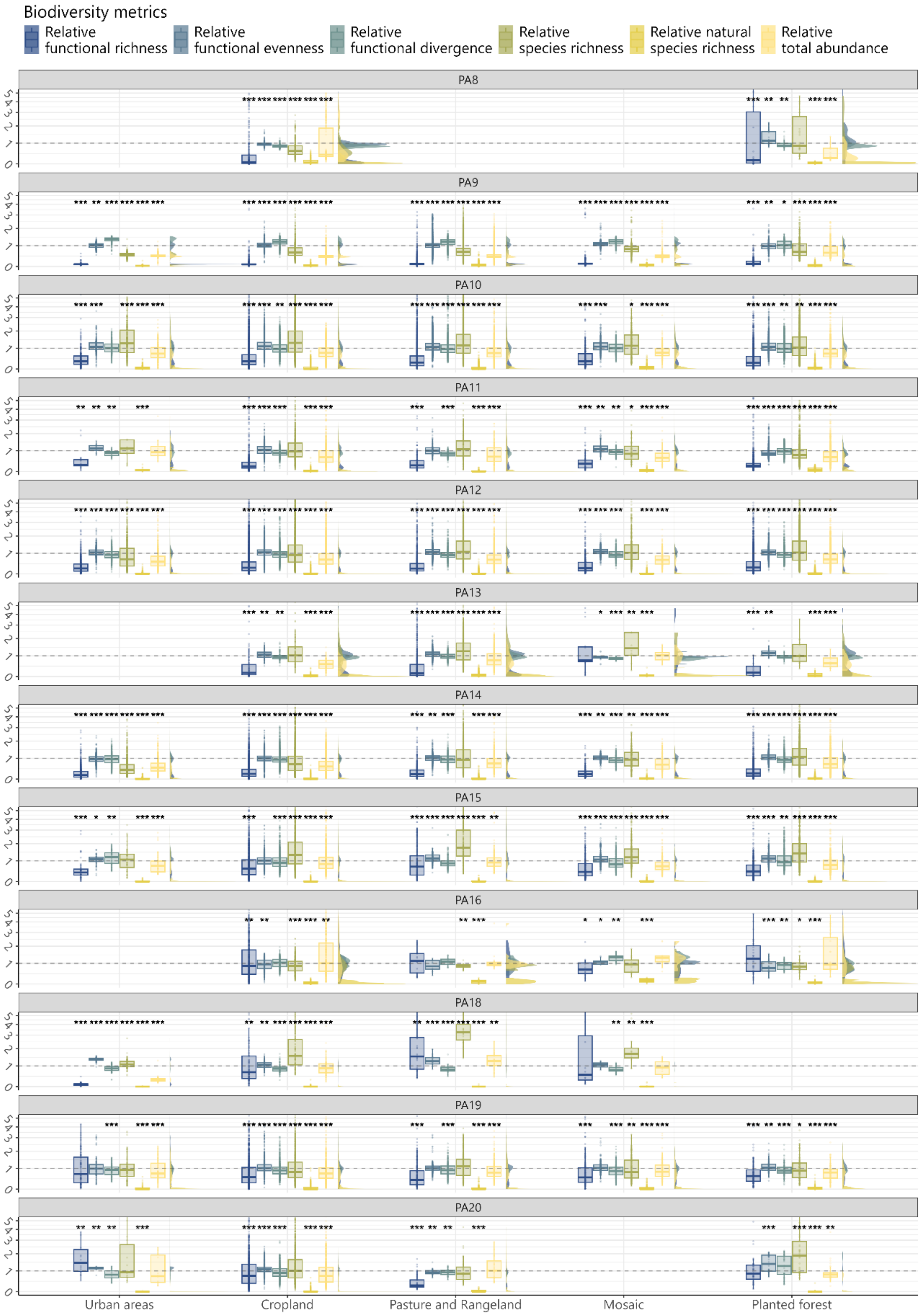


Figure 12.1 Relative values of the biodiversity metrics when vegetation plots with location uncertainty of ranking 3 are removed.

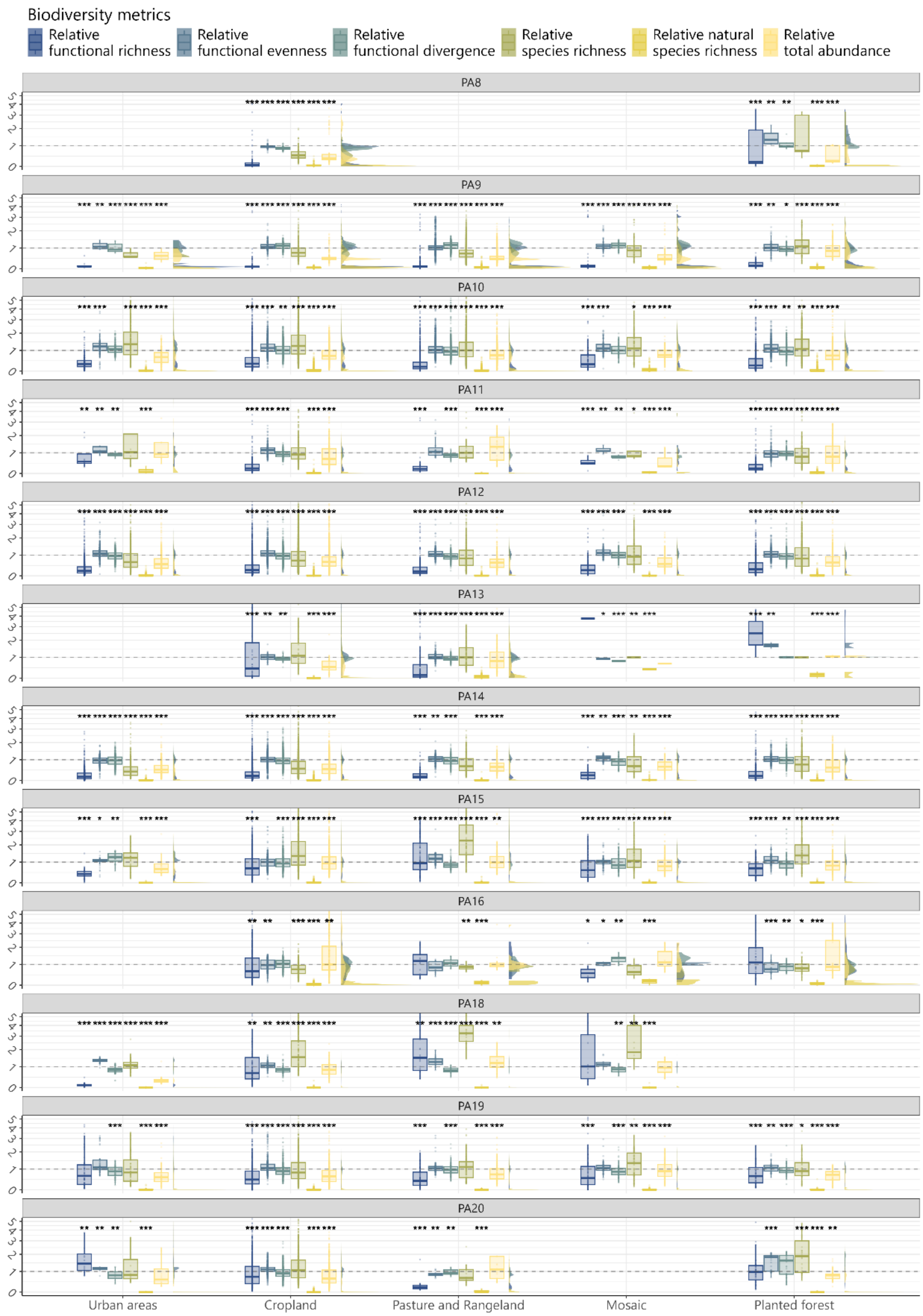


Figure 12.2 Relative values of the biodiversity metrics when pairs with distance between PC scores greater than 0.01 are removed.

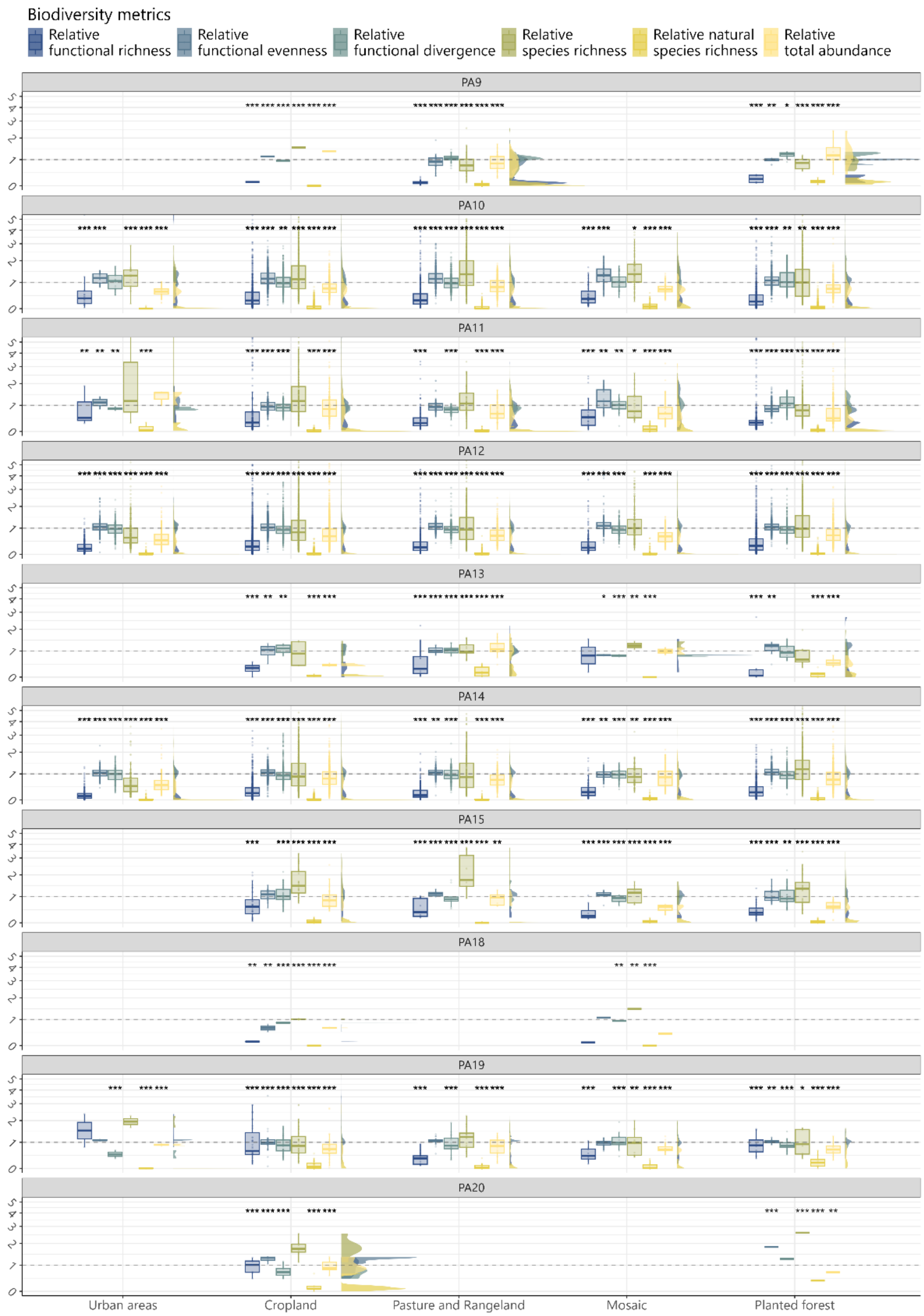


Figure.12.3 Relative values of the biodiversity metrics when pairs with anthropogenic vegetation plots assigned to naturalness level 2 are removed.

### S13 Absolute values of functional richness, functional evenness, functional divergence, species richness and total abundance and relative natural species richness

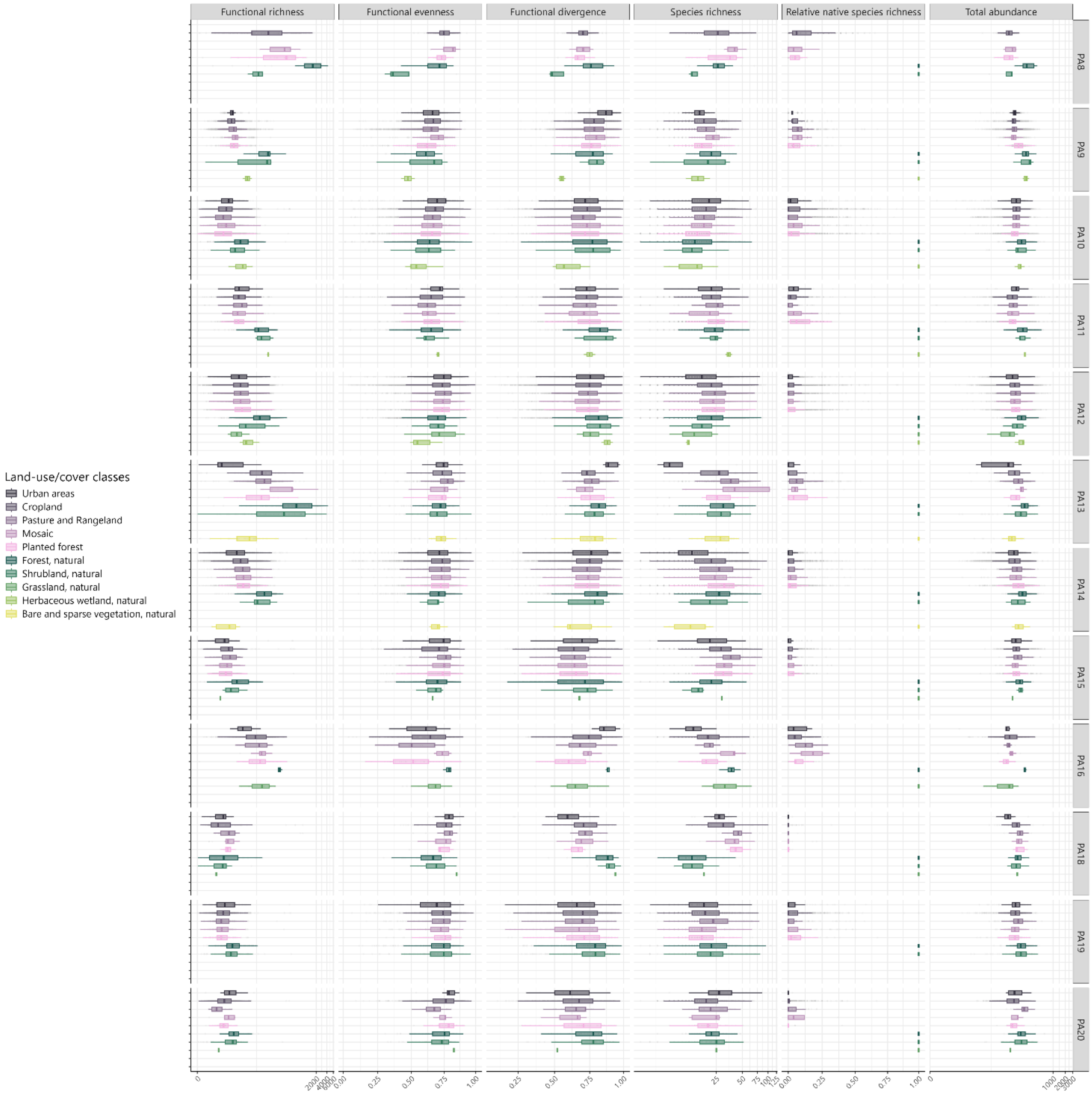


Figure S13.1 Values of biodiversity metrics per land-use/cover class and bioregion (the relative natural species richness is expressed in relative terms, as it depends on the paired natural habitat by definition).

#### S14 Relationship between species richness and functional richness

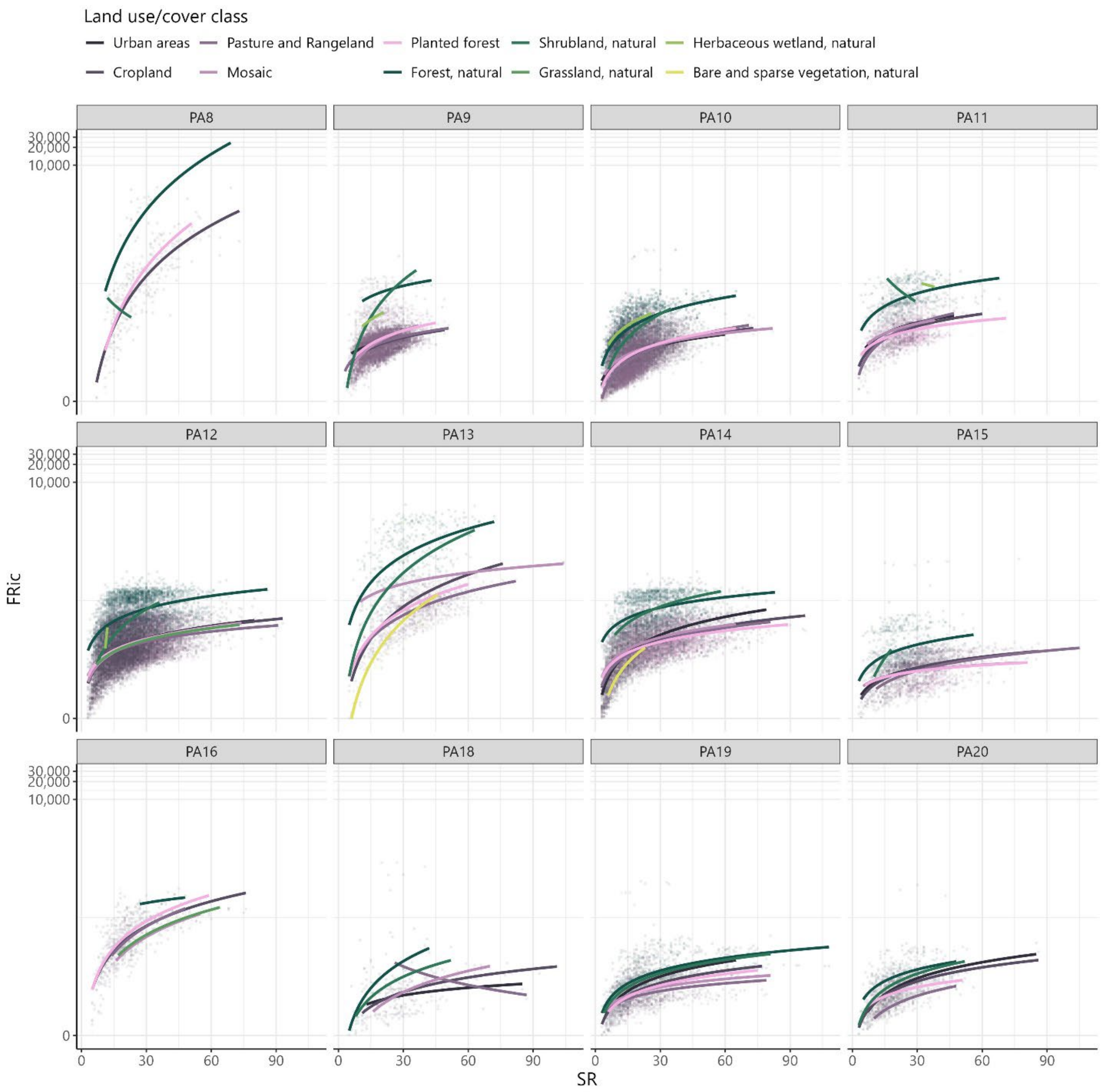


Figure S14.1 Scatterplot between species richness (SR) and functional richness (FRic).

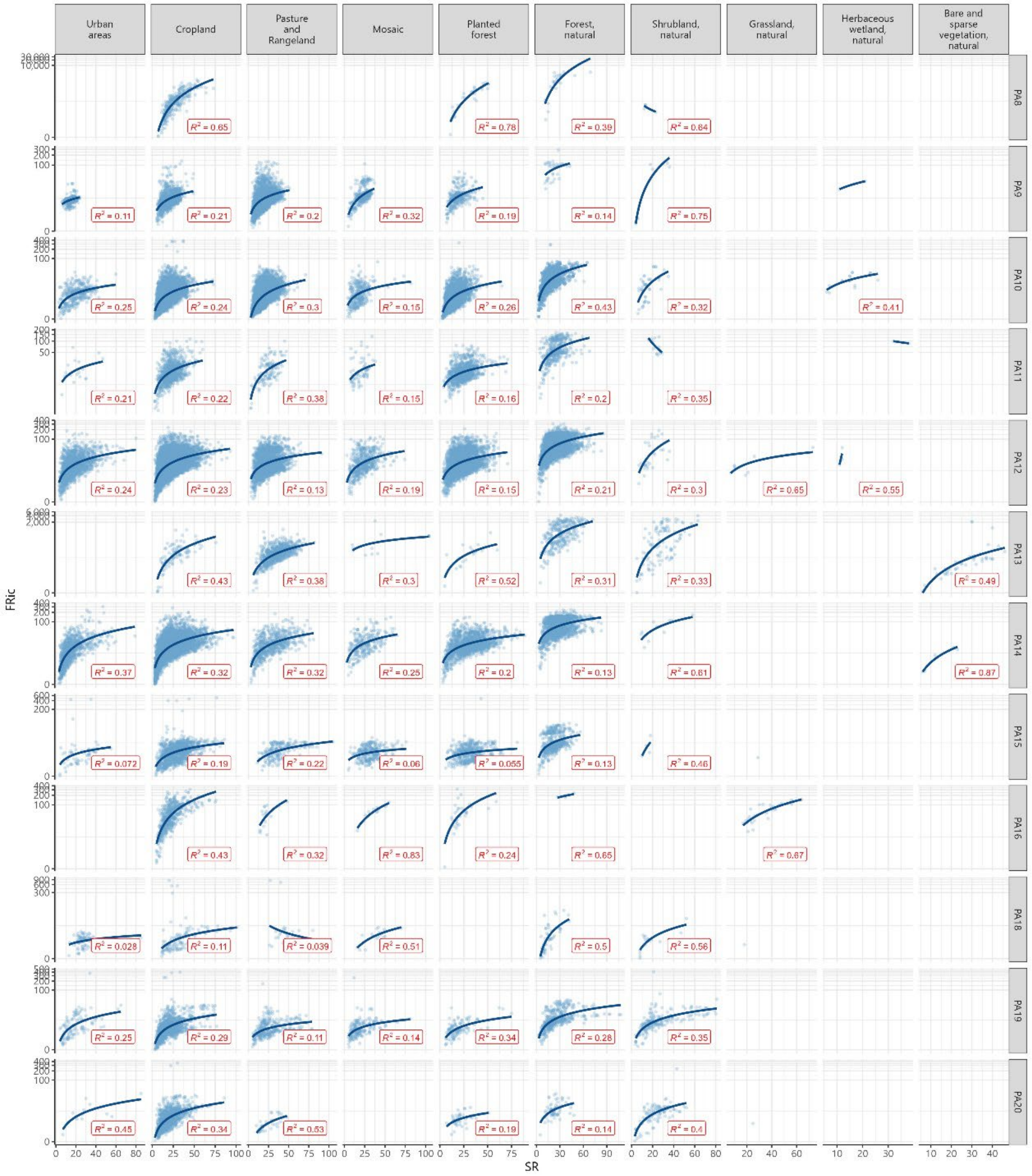


Figure S14.2 Scatterplot between species richness (SR) and functional richness (FRic) with $R^2$.

#### S15 Plot size and plot location uncertainty

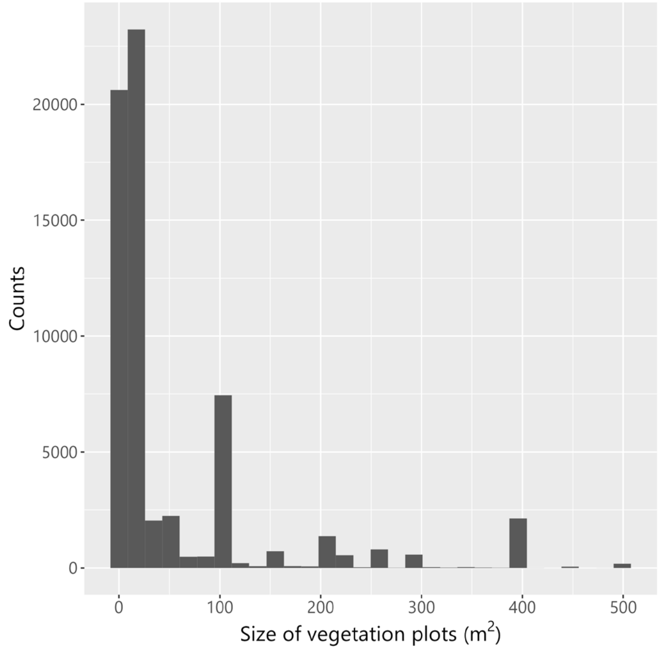


Figure S15.1 Distribution of the sample area of the vegetation plots considered in this study.

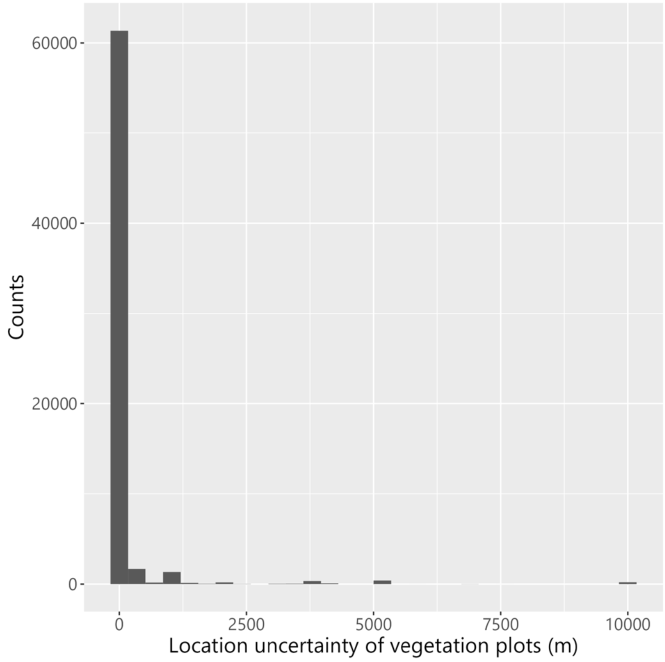


Figure S15.2 Distribution of the location uncertainty of the vegetation plots considered in this study.

#### S16 List of databases

.csv file containing the list of databases from EVA used to perform the analysis.

#### S17, S18 and S19 Numerical results

These files are available for review on Dryad at:

https://datadryad.org/stash/share/50nM9Rbd-qMSoiREeLir9CekoiFLG39bt3V5PiUdj4s

**S17**: .csv file (S17_abs_values_FD_SR_TotAb_RNSR_all-columns.csv) containing the numerical results of the functional and species diversity per vegetation plot, see the list of columns here below. This is the input file for the script main.R.

| Name | Description | Original from EVA/sPlot (*) or calculated/derived/ assigned in the study (**) |
|---|---|---|
| PlotObservationID | ID code of each vegetation plot | * |
| FRic | Functional richness | ** |
| FEve | Functional evenness | ** |
| FDiv | Functional divergence | ** |
| SR | Species Richness | ** |
| TotAb | Total abundance | ** |
| RNSR | Relative natural species richness | ** |
| broad_class | Land-use class or habitat | ** |
| distancePC | Distance of PC scores between the paired plots (see Pairing ID) | ** |
| Bioregion_code | Bioregion code | ** |
| PairingID | ID code of each pairing between vegetation plots | ** |
| Treatment | Anthropogenic land-use class (TRUE) or natural habitat (FALSE)? | ** |
| loc_uncertainty_ranking | Ranking of location uncertainty | ** |
| broad_class_PotNatVeg | Class of potential natural vegetation occurring at the coordinates of the plot | ** |
| Dataset | EVA dataset | * |
| GIVD.ID | EVA GIVD ID | * |
| Latitude | Latitute of the vegetation plot (rounded at 3 digits) | * |
| Longitude | Longitute of the vegetation plot (rounded at 3 digits) | * |
| Country | Country where the vegetation plot was sampled | * |
| Date.of.recording | Date of sampling | * |
| Location.uncertainty.m | Location uncertainty sampling (meters of the raius) | * |
| Releve.area | Sampling area (square meters) | * |
| Naturalness | Naturalness level from sPlot | * |
| Forest | Was the plot sampled in a forest (1 = yes, 0 = no)? | * |
| Shrubland | Was the plot sampled in a shrubland (1 = yes, 0 = no)? | * |
| Grassland | Was the plot sampled in a grassland (1 = yes, 0 = no)? | * |
| Wetland | Was the plot sampled in a wetland (1 = yes, 0 = no)? | * |
| Sparse.vegetation | Was the plot sampled in sparse vegetation (1 = yes, 0 = no)? | * |
| land_cover_code_ESA_CCI | ESA CCI land cover code extracted at the coordinates of the vegetation plot | ** |
| land_cover_code_HILDA | HILDA land cover code extracted at the coordinates of the vegetation plot | ** |
| label_ESA_CCI | ESA CCI land-cover category corresponding to the code | ** |
| label_HILDA | HILDA land-cover category corresponding to the code | ** |
| land_cover_code_NatureMap_ PotNatVeg | NatureMap land cover code of potential natural vegetation extracted at the coordinates of the vegetation plot | ** |
| label_NatureMap_PotNatVeg | NatureMap land-cover category corresponding to the code | ** |
| land_cover_code_GLOBIO | GLOBIO4 land cover code extracted at the coordinates of the vegetation plot | ** |
| label_GLOBIO | GLOBIO4 land-cover category corresponding to the code | ** |
| PC1 to PC23 | Results of the PCA on the soil and bioclimatic variables extracted at the coordinates of the vegetation plot | ** |

**S18**: .csv file (S18_relative_FD_SR_TotAb_RNSR.csv) containing the numerical results of the relative functional and species diversity per vegetation plot, see the list of columns here below. This is an output file of the script main.R.

| Name | Description | Original from EVA/sPlot (*) or calculated/derived/ assigned in the study (**) |
|---|---|---|
| PlotObservationID | ID code of each vegetation plot | * |
| FD_metric | Label of the functional or species diversity metric | ** |
| values | Value of the metric | ** |
| broad_class | Anthropogenic land-use class for which the relative value has been calculated | ** |
| distancePC | Distance of PC scores between the vegetation plot and its corresponding vegetation plot in the natural habitat | ** |
| Bioregion_code | Bioregion code | ** |
| PairingID | ID code of the pairing between each vegetation plot sampled in an anthropogenic land-use classes and the corresponding vegetation plot sampled in natural habitat | ** |
| loc_uncertainty_ranking | Ranking of location uncertainty | ** |
| pot_nat_veg | Class of potential natural vegetation occurring at the coordinates of the plot | ** |
| Wt_pvalue | p-value of BH-corrected Wilcoxon test for the combination of land-use class and bioregion to which the vegetation plot belongs | ** |
| n | Number of pairings in the combination of land-use class and bioregion to which the vegetation plot belongs | ** |
| Sign_Wt | Significance of the BH-corrected Wilcoxon test | ** |
| Latitude | Latitute of the vegetation plot (rounded at 3 digits) | * |
| Longitude | Longitute of the vegetation plot (rounded at 3 digits) | * |
| Country | Country where the vegetation plot was sampled | * |
| Releve.area | Sampling area (square meters) | * |
| Naturalness | Naturalness level from sPlot | * |

**S19**: .csv file (S19_aggr_abs_values_FD_SR_TotAb_RNSR.csv) containing the numerical results of the absolute values of functional and species diversity per anthropogenic land-use class/natural habitat, aggregated at bioregion level, see the list of columns here below (no original Eva/sPlot information, all derived). This is an output file of the script main.R.

| Name | Description |
|---|---|
| PlotObservationID | ID code of each vegetation plot |
| broad_class | Anthropogenic land-use class or natural habitat |
| Bioregion_code | Bioregion code |
| FD_metric | Label of the functional or species diversity metric |
| Mean | Mean of the absolute values of functional of species diversity per land-use class or natural habitat and bioregion |
| Median | Median of the absolute values of functional of species diversity per land-use class or natural habitat and bioregion |
| Stand_dev | Standard deviation of the absolute values of functional of species diversity per land-use class or natural habitat and bioregion |
| n | Number of pairings in the combination of anthropogenic land-use class and bioregion |

**S20**: .csv file (S20_aggr_relative_FD_SR_TotAb_RNSR.csv) containing the numerical results of the relative values of functional and species diversity per anthropogenic land-use class, aggregated at bioregion level, see the list of columns here below (no original Eva/sPlot information, all derived). This is an output file of the script main.R.

| Name | Description |
|---|---|
| PlotObservationID | ID code of each vegetation plot |
| broad_class | Anthropogenic land-use class |
| Bioregion_code | Bioregion code |
| FD_metric | Label of the functional or species diversity metric |
| Sign_Wt | Significance of the BH-corrected Wilcoxon test |
| Mean | Mean of the relative values of functional of species diversity per land-use class or natural habitat and bioregion |
| Median | Median of the relative values of functional of species diversity per land-use class or natural habitat and bioregion |
| Stand_dev | Standard deviation of the relative values of functional of species diversity per land-use class or natural habitat and bioregion |
| n | Number of pairings in the combination of anthropogenic land-use class and bioregion |
